# Untargeted metabolomic analyses reveal the diversity and plasticity of the specialized metabolome in seeds of different *Camelina sativa* genotypes

**DOI:** 10.1101/2021.01.18.427130

**Authors:** Stéphanie Boutet, Léa Barreda, François Perreau, Jean-Chrisologue Totozafy, Caroline Mauve, Bertrand Gakière, Etienne Delannoy, Marie-Laure Martin-Magniette, Andrea Monti, Loïc Lepiniec, Federica Zanetti, Massimiliano Corso

**Author notes:** Correspondence to: Massimiliano Corso. Equal contribution (Stéphanie Boutet, Léa Barreda). E-mails: SB.

## Abstract

Despite the essential role of Specialized Metabolites (SMs) in the interaction of plants with the environment, studying the ability of crop seeds to produce these protective compounds has been neglected. Furthermore, seeds produce a myriad of SMs providing an interesting model to investigate their diversity and plasticity. *Camelina sativa* gained a lot of interest in the past few years as rustic oil seed crop. A characterization of seed SM landscapes in six camelina genotypes grown in the field and harvested during five growing seasons has been undertaken in this work. This allowed a comprehensive annotation of seed SMs combining analyses that cluster SMs based on their chemical structures and co-accumulation patterns. These data showed broad effects of the environment on the stimulation of the seed-specialized metabolome. Among well annotated compounds, flavonols were identified as the metabolic class characterised by high plasticity, revealing significant variable accumulation according to the year and/or the genotype. Lastly, a deeper characterisation of primary metabolites and lipids in two selected genotypes has been performed. We showed that, in addition to flavonols, alkaloids and glucosinolates displayed a higher phenotypic plasticity with respect to most of the primary metabolites, including some sugars and major storage compounds such as fatty acids, proteins and most lipid classes (e.g. DAG, TAG), but similar plasticity compared to free aminoacids and carboxylic acids. This work highlighted major and unexplored effects of the environment on the seed specialized metabolome demonstrating that seeds exhibit a dynamic and plastic metabolism, with an impact on seed quality.

**Significance statement:** Seeds produce a myriad of Specialized Metabolites (SMs) with an essential role in the interaction of plants with the environment. We characterized SM landscapes, primary metabolites and lipid composition in the seeds of camelina genotypes grown in the open field in five consecutive growing seasons. Our results showed the predominant effect of the environment on the regulation of the seed - specialized metabolome, with a potential impact on seed quality of camelina that may also occur in other oilseed crops.

## INTRODUCTION

Plants produce large amounts of metabolites with different functions ranging from environmental adaptation to developmental controls (Erb and Kliebenstein, 2020; Fang *et al*., 2019). Some of these metabolites, namely specialized (or secondary) metabolites (SMs), are usually synthetized in small concentrations, and have a limited distribution to specific plant families, genera or species (Tissier *et al*., 2015; Bhatla, 2018; Pichersky and Lewinsohn, 2011; Arimura and Maffei, 2016). They are commonly classified in three major groups according to their chemical structure: terpenoids, nitrogen containing molecules, and phenolic compounds (or phenylpropanoids) (Jacobowitz and Weng, 2020). In addition, many lipids are also characterized by huge diversity among plant species and many of them can be considered as SMs (Matsuzawa *et al*., 2021; Tsugawa *et al*., 2020). Hence, each class of metabolites comprises a huge diversity of structures leading to a wide range of biological activities (Wang *et al*., 2019; Kosmacz *et al*., 2020).

In plants, terpenoids are the most structurally divers group of annotated SMs, with more than 25000 compounds identified that derived from the assembly of 5-C isoprene skeletons (terpenes) (Tissier *et al*., 2015). A second group of SMs, referred as nitrogen-containing compounds, includes alkaloids, glucosinolates, cyanogenic glycosides and some non-protein amino acids (Wang *et al*., 2019; Ziegler and Facchini, 2008; Halkier and Gershenzon, 2006). Among them, alkaloids are the most represented, with more than 15000 compounds annotated (Bhatla, 2018). Finally, phenolic compounds (or phenylpropanoids) are molecules containing one or more hydroxylated aromatic rings, and are synthetized via the malonate and shikimate pathways (Vogt, 2009; Tohge *et al*., 2017; Corso *et al*., 2020). Plants produce more than 10000 phenolic compounds, which range from simple single aromatic rings such as benzoic acid or cinnamic acid to more complex molecules as flavan-3-ol polymers (also termed proanthocyanidins, PAs) (Corso *et al*., 2020; Lepiniec *et al*., 2006). Flavonoids are certainly the largest group of phenylpropanoids, with more than 9000 representatives (Tohge *et al*., 2017; Corso *et al*., 2020). They have a C6-C3-C6 skeleton, characterized by two benzene rings fused by a heterocyclic pyran ring. Flavonoid diversity, biological activity and transport are strongly affected by the degree of glycosylation, hydroxylation and methylation patterns of the benzene rings (Tohge *et al*., 2020; Sudheeran *et al*., 2020; Tohge *et al*., 2017; Wang *et al*., 2019).

Specialized metabolites play crucial physiological and ecological roles throughout the plant life cycle. They protect plants against abiotic stresses such as drought, high or low temperatures, toxic metals and UV, and can constitute effective defences against biotic stresses such as pathogens, herbivores or plant competitors (Delgoda and Murray, 2017; Corso *et al*., 2020; Lacchini and Goossens, 2020; Kosmacz *et al*., 2020; Lepiniec *et al*., 2006; Li *et al*., 2020; Tohge *et al*., 2016). For example, SMs can act as phytoalexins in response to pathogens, antifeedants against herbivory or allelopathic compounds to stunt the germination/growth of other plants (Corso *et al*., 2020; Corso *et al*., 2021). These compounds are also involved in plant nutrition and/or reproduction, they play a role in plant metal homeostasis and in plant-microbes interaction, as in the case of root flavones affecting composition rhizosphere microbial community to improve maize performance upon nitrogen deprivation (Corso and de la Torre, 2020; Corso *et al*., 2020; Croteau, R., Kutchan, T.M. and Lewis, 2000; Maeda, 2019; Arimura and Maffei, 2016; Korenblum *et al*., 2020; Yu *et al*., 2021). Given the key role of SM in plant-environment interaction, metabolomics is considered the omics technique that is closest to the phenotype (Guijas *et al*., 2018). In fact, SMs give a substantial contribution to plant phenotypic plasticity, defined as the ability of a genotype to express different phenotypes (in this case the metabolites) according to environmental conditions (Fox *et al*., 2019; Bonamour *et al*., 2018; Diouf *et al*., 2020; Bradshaw, 1965; Grenier *et al*., 2016).

Seeds constitute (directly or indirectly) a major component of the human diet, providing proteins, oil, and starch for both human and animal nutrition. They present a wide range of SMs that are important for their physiological, agronomic, nutritional and industrial qualities (Corso *et al*., 2020; Lepiniec *et al*., 2006; Corso *et al*., 2021). A large diversity of SMs has been characterized in seeds of several plant species, including *Brassicaceae* (Routaboul *et al*., 2006; Quéro *et al*., 2016; Corso *et al*., 2020; Lepiniec *et al*., 2006). They show remarkable levels and wide distribution of phenylpropanoids, flavonoids in particular (Routaboul *et al*., 2006; Routaboul *et al*., 2012; Alseekh, Ofner, *et al*., 2020; Auger *et al*., 2010). Nevertheless, we are far from an exhaustive characterization of seed-specialized metabolite diversity and of the impact of the environment on this diversity.

Camelina [*Camelina sativa* (L.) Crantz] is a *Brassicaceae* species that has gained a lot of interest in the past few years (Zanetti *et al*., 2021). This crop is highly tolerant to biotic and abiotic stresses. Camelina seeds are characterized by unique oil composition (e.g. high level of polyunsaturated fatty acids, PUFA, and in particular 18:3 fatty acids), and high oil content. It reaches adequate yields under low-input management and in limiting environments (Zanetti *et al*., 2017; Faure and Tepfer, 2016; Yuan and Li, 2020). Nevertheless, further studies are needed to better valorise camelina seed co-products after oil extraction, including SMs such as polyphenols and carotenoids present in high amounts (Quéro *et al*., 2016). Only a few data are available on camelina SM diversity and no information is available on their plasticity and induction in response to environmental conditions.

In this work, we characterized the diversity of specialized metabolites in the seeds of six camelina genotypes (MIDAS, OMEGA, WUR, 787-08, 789-02 and 887) grown in the open field and harvested during five consecutive growing seasons (2015-2019). Using molecular network analyses, which gather metabolites based on their structures, combined with metabolite co-accumulation networks, we extended the characterization and pathway information of the main metabolic classes in camelina seeds. Among the main seed SM classes, we identified those influenced by the environment and/or the genotype. Moreover, we characterized the impact of the environment on the phenotypic plasticity of primary and specialized metabolites in camelina seeds.

## RESULTS

### Plant development and meteorological data

In the multi-year experiment (2015-2019) conducted on six camelina genotypes grown in Italy (Bologna), the plants were able to reach maturity in about 100 days (Figure 1a). The average minimum (T_min_) and maximum (T_max_) temperatures (Figure 1b) measured from 50% flowering to seed maturity (Figure 1c), a particularly important phase for seed quality, were quite variable in the years studied. This particular developmental phase has been reported as the most important in camelina seed development and filling (Walia *et al*., 2018; Righini *et al*., 2019). In detail, the average T_min_ ranged from 13.5 °C in 2017 to 16.6 °C in 2019, while the average T_max_ ranged from 25.6 °C in 2016 to 29.3 °C in 2019. The cumulative precipitation in the same growth phase (from 50% flowering to maturity) showed even higher variability among the five years studied (Figure 1b). Specifically, 2017 had the lowest precipitation (52.2 mm), whereas 2016 showed around two- to three-fold higher cumulative precipitation (161.6 mm) compared with the other years (Figure 1b). Besides, in 2016, a hailstorm occurred before harvest (Zanetti *et al*., 2017). Considering these variations, we will refer to the multiple factors underlying differences among growing seasons as the ‘environment’.

**Figure 1.**
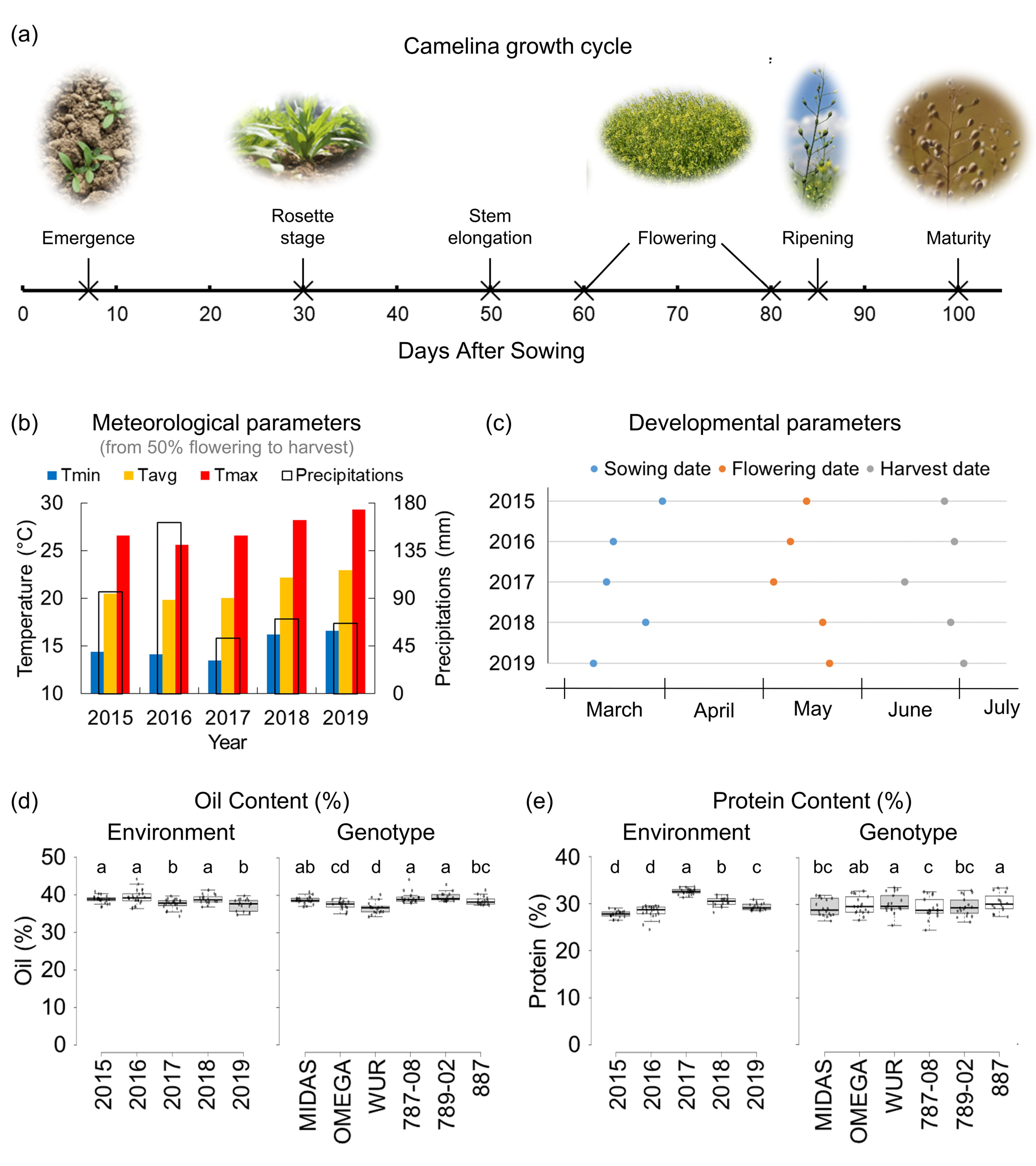
Growth cycle of camelina, meteorological data and seed quality parameters. (a) Developmental cycle of camelina plants in Bologna (Italy). The average phenological stages and days after sowing (DAS) are indicated. (b) Meteorological parameters obtained from 50% flowering to seed maturity. Minimum (Tmin), average (Tavg) and maximum (Tmax) temperatures (°C), and amount of precipitation (mm) are shown. (c) Sowing, flowering and harvest dates are indicated for each year studied (2015 to 2019). (d, e) Total seed oil (d) and protein (e) contents are shown for each year and genotype. Vertical bars = standard deviations. Different letters indicate statistically significant differences (*P* ≤ 0.05, Tukey’s range test).

### Plant phenology and seed quality parameters

Environmental conditions had a significant influence on camelina development across years (Figure 1c). In particular, sowing took place between mid-March to early April, and consequently flowering occurred between early and late May, while seed maturity was reached between mid-June and early-July. Seed oil content (%) showed low variations among genotypes and years (Tukey’s range test; *P*≤0.05). WUR was the genotype with the lowest average seed oil content (36.7 %), while 787-08 and 789-02 (both 39.4 %) reached the highest values. Concerning years, 2016 (39.3 %) and 2019 (37.4 %) reported the highest and the lowest seed oil content, respectively, compared with the other growing seasons. Seed protein content (%) was also significantly, albeit modestly, influenced by years and genotypes. Among genotypes, 787-08 (29.2 %) had the lowest protein content, and 887 and WUR the highest (30.1 %). Concerning years, 2017 was the one reporting the highest seed protein content, while 2015 and 2016 had lowest protein content. It is worth mentioning that the coefficient of variation for seed oil content and seed protein content was only 4.2% and 6.7% respectively, highlighting the limited variability for these qualitative traits in camelina seeds (Figures 1d, e).

### The metabolomic landscape of camelina seeds

Untargeted SM profiling revealed a large diversity in camelina seeds (Figure 2; Table S1). Untargeted metabolomic data in ElectroSpray Ionization in positive (ESI+) and negative (ESI-) modes allowed the detection of 1595 peaks of putative known and unknown metabolites (i.e. metabolic features).

**Figure 2.**
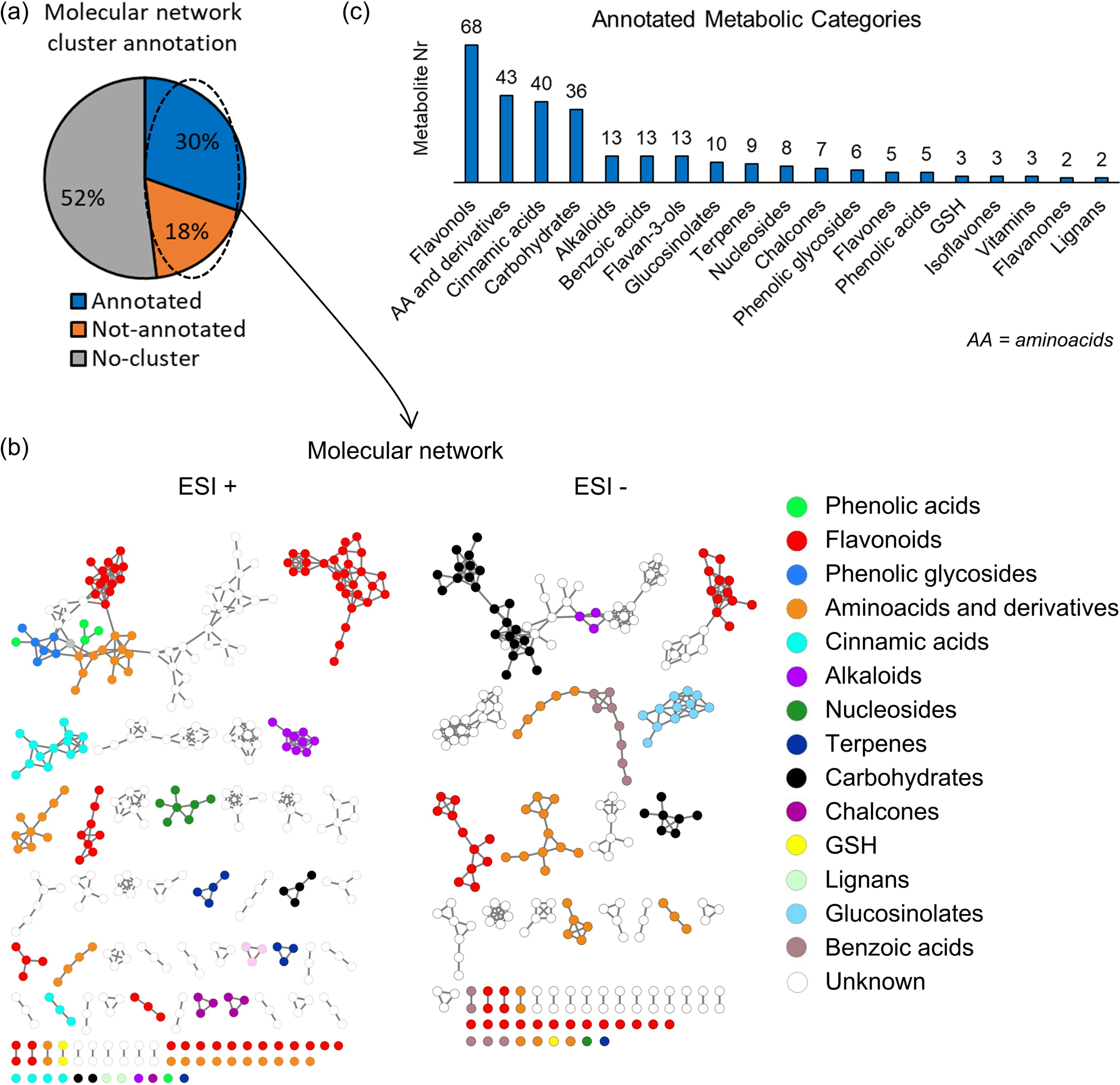
Molecular Networks and annotation of the seed-specialized metabolites (a) The molecular networks are shown for metabolite analyses carried out in positive (ESI+) and negative (ESI-) ionization modes. Different colours correspond to different metabolic classes. Metabolites are grouped based on their chemical structures. Cosine similarity scores of 0.8 and 0.75 were used for ESI+ and ESI-, respectively. (b) Percentage of specialized metabolites present in a molecular network with or without an annotation. (c) Histogram showing the annotated/known metabolic categories identified in camelina seeds. The number of metabolites that belong to each category is indicated in the figure.

The average intensity of SMs among all samples (Table S2) was calculated and identified the top-10 with higher average intensity (Figure S1a). Among them four were flavonoids (Rutin, Quercetin-3- (Glucose [G] - Rhamnose [R] - Pentose [P]), Isorhamnetin-3-(G - R) and Catechin-3-G), one cinnamic acid (Sinapine), and one aminoacid (γ-glutamyl leucine). In addition, four compounds had no annotation in homemade or public libraries (n8, n14, p29, p117; Figure S1b). The putative structure of these metabolites with no annotation was determined using SIRIUS 4.0 (Dührkop *et al*., 2019), that propose molecular structures using LC-MS/MS data (see methods). The proposed structures of n8 and n14 belong to the glucosinolate class, more precisely n8 was annotated as Glucocamelinin and n14 as Glucoarabin. p29 was putatively annotated as a sugar while metabolite p117 was annotated as an aminoacid.

LC-MS/MS data were used to build a molecular network with MetGem software (Figure 2a, b; Table S1). This method allows clustering of compounds with high degree of structural similarity according to their MS^2^ spectra, and therefore identifying clusters of unknown metabolites using the MS/MS information for annotated compounds (Olivon *et al*., 2018). The analysis of camelina seed metabolites using public databases for annotation and internal standard, coupled to molecular network, allowed the assignment of 30% of the detected molecules to a metabolic category. Conversely, 18% of the metabolites were included in clusters that could not be associated to a metabolic class, and 52 % of the metabolites were not comprised in a cluster and not annotated (Figure 2a). Overall, SMs identified in camelina seeds have been assigned to sixteen putative metabolic categories: flavonoids (flavonols, flavan-3-ols, flavones, flavanones and isoflavones), aminoacids and derivatives, cinnamic acids, carbohydrates, alkaloids, benzoic acids, glucosinolates, terpenes, nucleosides, chalcones, phenolic glycosides, phenolic acids, glutathione (GSH) and lignans (Figure 2c). Flavonols were the most represented metabolic category (68), followed by aminoacids and derivatives (43), cinnamic acids (40) and sugars and carbohydrates (36). A relatively lower number of metabolites were annotated as alkaloids (13), flavan-3-ols (13), benzoic acids (13) and glucosinolates (10). The other metabolic categories presented less than 10 metabolites.

### Genetic and environmental regulations of camelina seed specialized metabolome

The ESI+ and ESI-relative metabolite intensities data were analysed by a linear model to evaluate the genotype and year effects as well as their interactions. After filtering, statistical analyses were performed with 1481 out of the 1595 metabolic features detected (see experimental procedures). Pairwise comparisons were carried out between the different years (*e.g.* 2015 vs. 2016 on average for all genotypes), genotypes (*e.g.* OMEGA vs. MIDAS on average for all years) and genotype X environment (*e.g.* [OMEGA-2015 vs. OMEGA-2016] compared with [MIDAS-2015 vs. MIDAS-2016]). From this point on Differentially Accumulated Metabolic features (DAMf) will be used to indicate the number of metabolic features showing significantly different levels (induced or repressed) between two samples or group of samples/factors. Statistical analyses highlighted 1111 and 1378 DAMf in response to the genotype and the environment (*i.e.* the year), respectively (Figure 3a). Furthermore, 1452 DAMf were affected by genotype X environment interaction. Thus, statistical analyses revealed that the environment was the most important factor influencing metabolite accumulation and affected 93% of the analysed camelina seed-specialized metabolome. To measure the specific impact of each year and genotype, DAMf were identified for each comparison (Figure 3b, Figure S2). Overall, the year 2016 showed the largest number of cumulated DAMf, with 2494 induced and 660 reduced metabolites compared to the other years among all comparisons (Figure 3b, Figure S2). Concerning genotypes, OMEGA showed the largest number of cumulated DAMf, with 1690 and 402 induced and reduced accumulation of SMs, respectively, compared with the others (Figure 3b, Figure S2).

**Figure 3.**
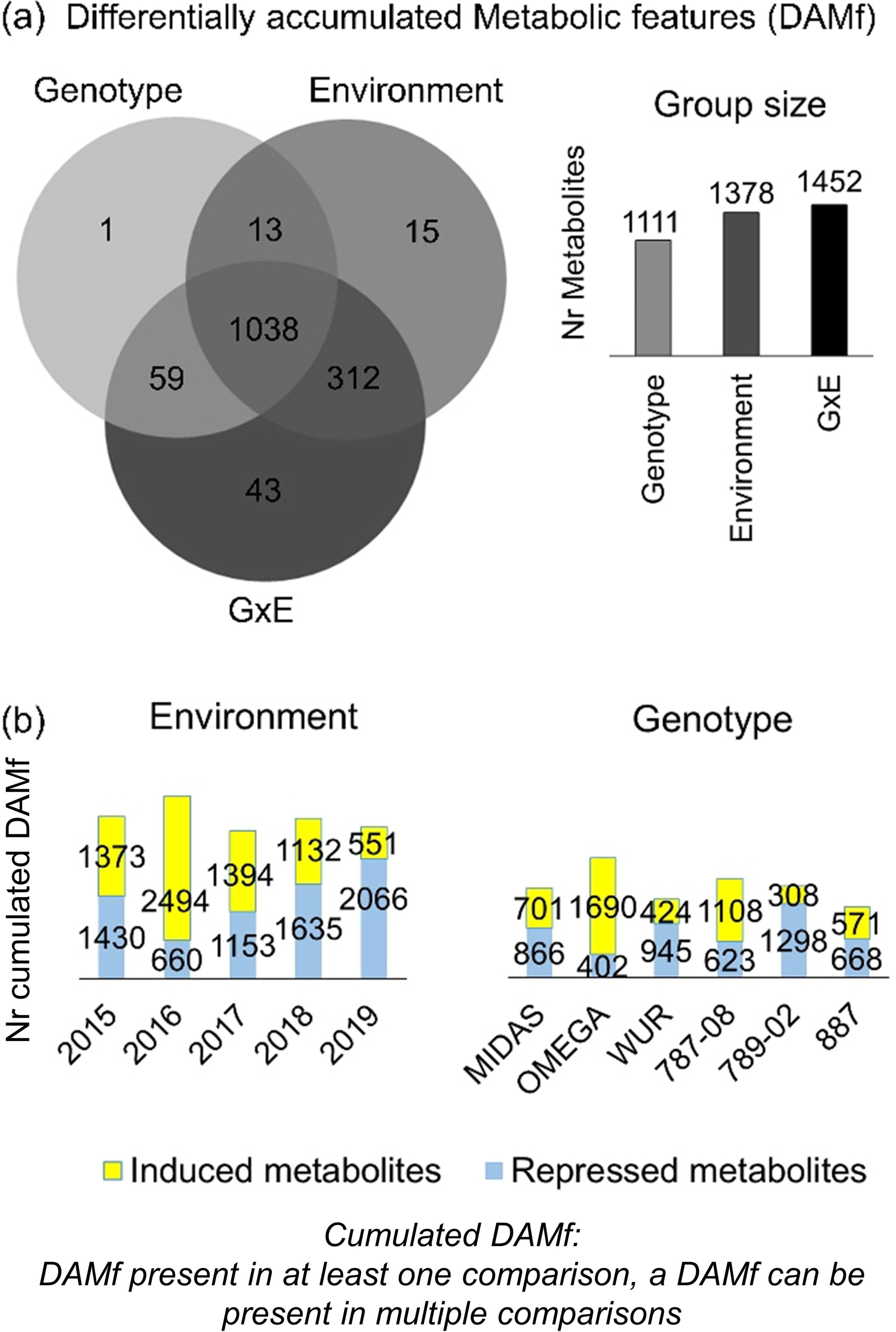
Genetic and environmental impact on the accumulation of seed-specialized metabolites. (a) Venn diagram and histogram indicating the number of Differentially Accumulated Metabolites (DAMf) according to the Environment, Genotype and Genotype x Environment (GxE) factors. (b) Number of induced (yellow) and repressed (blue) DAMf for each year and genotype. The DAM numbers are cumulative and were calculated considering all the statistical comparisons for each year and/or genotype. (c) Hierarchical clustering analysis using the amounts of metabolites. The results are shown separately for years and genotypes.

### Identification and accumulation profiles of metabolites affected by environmental and genotype variations

The analyses highlighted a strong and broad effect of the environment on SM diversity and accumulation, and the majority of DAMf that were in common between the environment and the genotype (G-E). In this light, the G-E DAMf (1051 metabolic features) were considered for further analyses (Figure 3a). Using this subset of DAMf, we performed an enrichment analysis using the metabolic categories highlighted by the molecular network analysis (Table S1, column ‘Metabolic category’). Not-annotated metabolites clustering together in the molecular network analysis (Figure 2b) and therefore having similar chemical structures, were considered as a distinct metabolic category (see methods for more information). Among the DAMf common to genotypes and environments, flavonols (47 DAMf), cinnamic acids (35 DAMf), and glucosinolates (10 DAMf) were found to be enriched compared with the metabolite reference set library (see methods for further information; Figure 4a). In addition, two unknown categories, namely p_2_a (20 DAMf) and p_2_c (9 DAMf) were also enriched (Figure 4a). The heatmaps built using the average metabolite intensity for each category showed that the environmental conditions of 2016 induced higher average metabolite accumulation compared with the other years for the enriched metabolic categories, except for flavonols and, to a lesser extent, to not annotated category p_2_c (Figure 4b). Conversely, 2018 and 2019 were the years during which cinnamic acids, glucosinolates, p_2_a and p_2_c metabolic categories displayed the lowest accumulation levels. In addition, the genotypes OMEGA and 787-08 showed higher amounts of metabolites for most of the categories. These results confirmed the strong effect of the environment, particularly in 2016, on the accumulation of SMs. Interestingly, flavonols presented an accumulation pattern that significantly differed from the other metabolite categories (Figure 4b).

**Figure 4.**
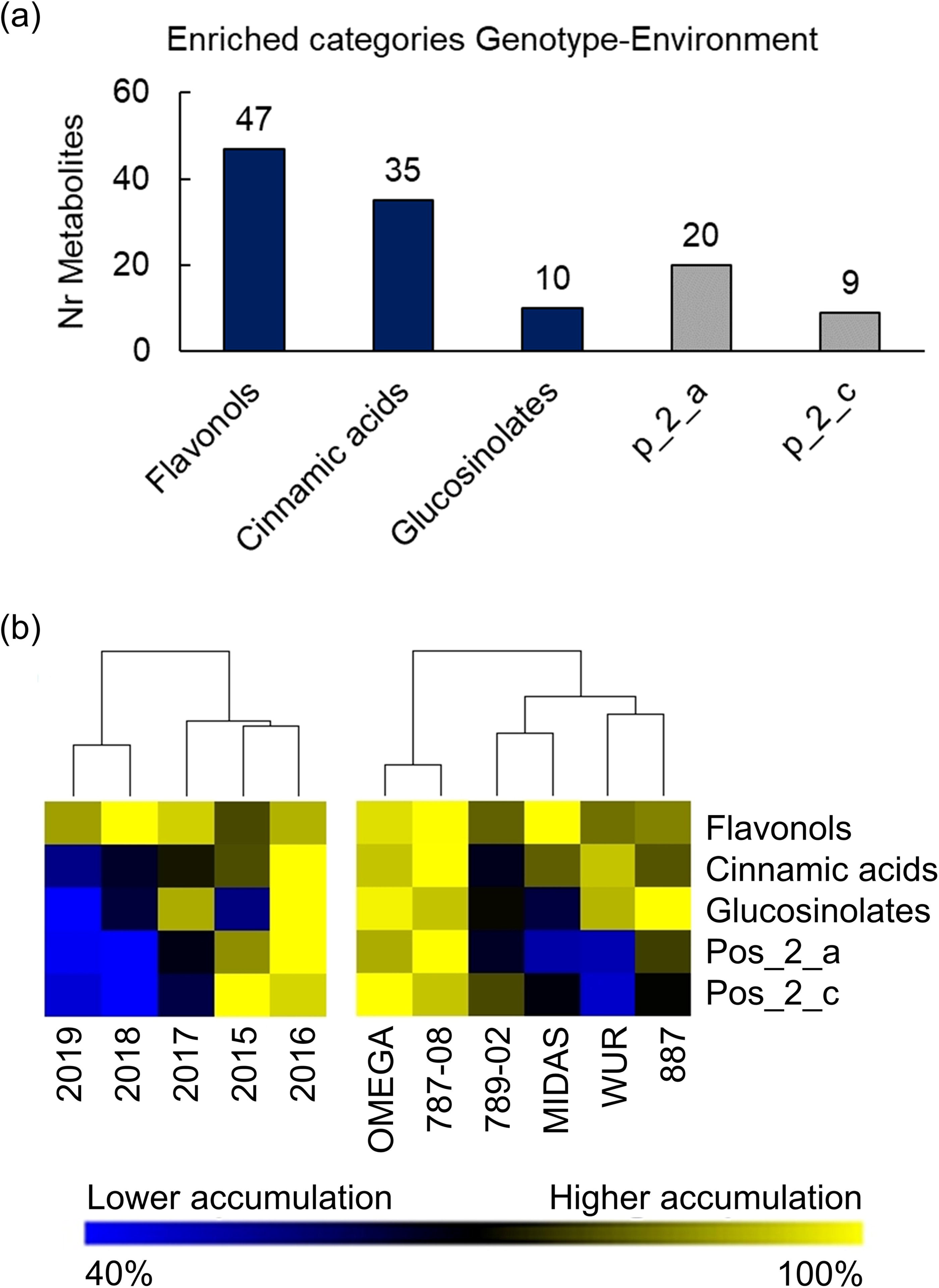
Enriched metabolic categories (a) Histograms showing the annotated and not-annotated enriched metabolic categories using DAM affected by the genotype (G) and the environment (E). The number of metabolites that belong to each category is indicated. (b) Heat maps showing the average metabolite accumulation separately for years and genotypes for each enriched metabolic category. The accumulation values were calculated as a percentage related to the sample showing the highest accumulation value for each metabolite (100% and 40% for yellow and blue colours, respectively) within years or genotypes.

### Identification of important metabolite hubs and their environmental induction

The results obtained so far highlighted large variations according to the environment and/or the genotype of several metabolic categories. Nevertheless, it remained unclear how genotypes or the environmental conditions affected the accumulation of specific metabolites or their decorations (e.g. glycosylated / acylated / methylated quercetin or kaempferol). To establish a link between metabolite structures (and therefore their putative/potential biological activity) and environmental conditions or genotypes, a co-accumulation network analysis was performed based on correlation among metabolite intensity (Figure 5; *r* > 0.7) using metabolomic data extracted for the enriched metabolic classes (Figures 6 and S4). Co-accumulated metabolites obtained from the metabolite correlation network were visualised in boxplots for each cluster. In comparison with the other enriched categories, which were grouped in only one cluster, flavonols were splitted into several clusters, highlighting more diversified accumulation profiles in response to year and/or genotype (Figure 6). A metabolic signature was identified for several flavonol clusters identified by co-accumulation analyses, and specific metabolite decorations were associated to each year (i.e. to different environmental conditions). Specifically, rutin (Quercetin [Q])-3-(G-R)), Kaempferol [K]-3-(G-R), Isorhamnetin [Iso]-3-(G-R), K-3-(G-R), Q-3-(G-R-P), Iso-3-(G-R-P) and Q-3-(G-R-R) (Figure 7), belong to a co-accumulation cluster that showed slightly lower accumulation in 2015 (cluster ‘a’). Other flavonol clusters showed a much clearer metabolite accumulation behaviour. In particular, acetylated and malonylated mono- or di-glycosylated quercetin and isorhamnetin belong to cluster ‘b’, characterized by metabolites showing markedly high accumulation in 2016, and in 789-02 and OMEGA genotypes. Cluster ‘c’, which is characterized by 12-times higher accumulation of metabolites in 2019 compared to other years, includes K-3-(G-Malonyl), K-(OH-3-MethylGlutaryl)-G and Q-3-(G-Galloyl). Q-3-(R-G-G) (cluster ‘d’) and diglycosylated quercetin bounded to feruloyl (cluster ‘f’) were markedly more accumulated in 2016, but their accumulation patterns differ according to the genotypes. Finally, metabolites belonging to cluster ‘e’, which are characterized by higher accumulation in 2018 and in MIDAS and 787-08, were tri-glycosylated quercetin and flavonols with coumaroyl, feruloyl or sinapoyl alcohol groups (Figure 7).

**Figure 5.**
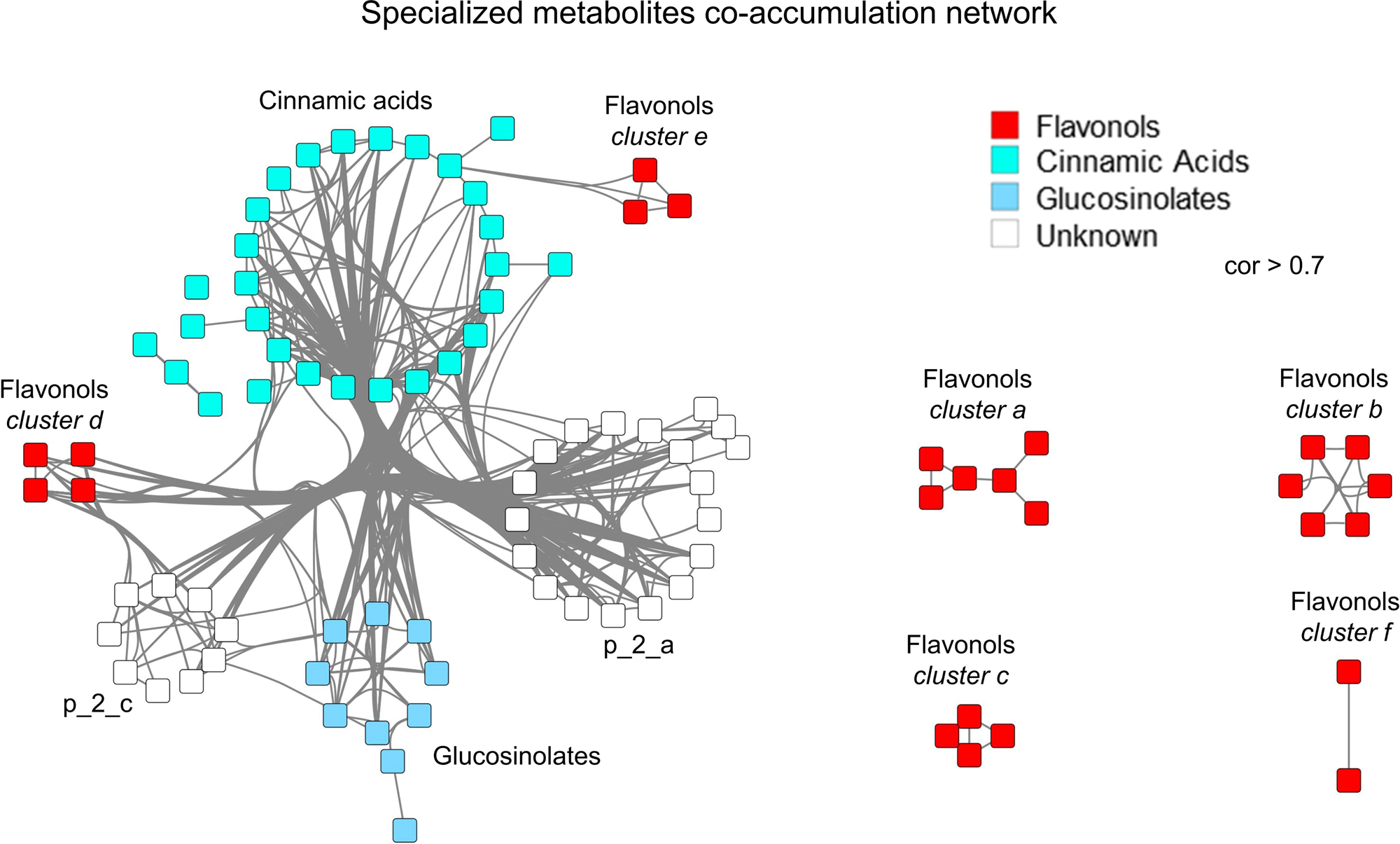
Specialized metabolites co-accumulation networks. The accumulation patterns were used to calculate correlations (r > 0.7) among metabolites and define the clusters. Only the metabolites belonging to enriched categories were considered in this analysis. The metabolites from each enriched category are highlighted with different colours.

**Figure 6.**
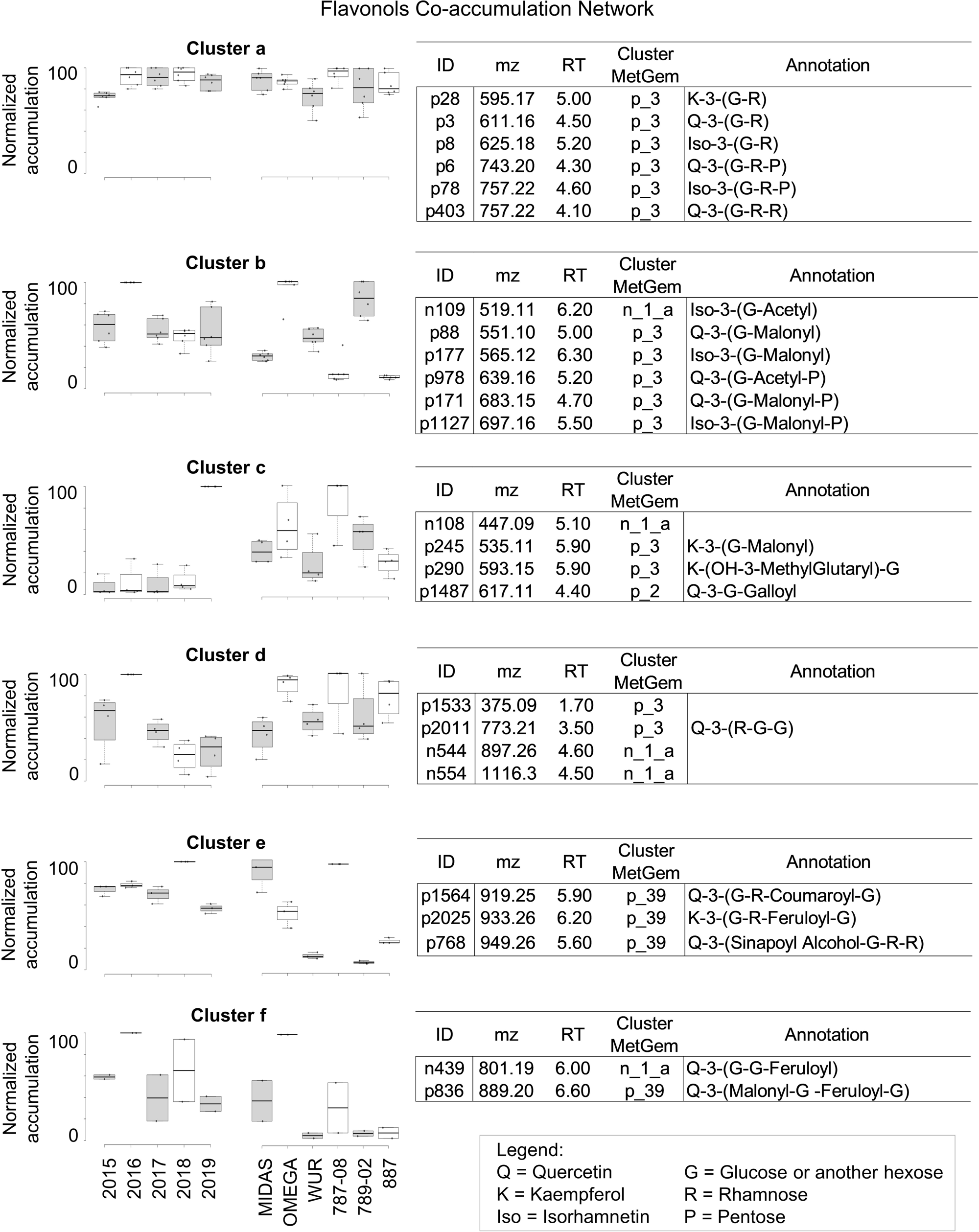
Flavonol co-accumulation network. Clusters and accumulation patterns are shown for the flavonols present in co-accumulation networks. The boxplots represent the accumulation patterns for each metabolite in all years and genotypes. The *m/z*, retention time, MetGem cluster and annotations are also shown for each metabolite. The accumulation values (boxplots) were normalized and calculated as a percentage related to the sample showing the highest accumulation value for each metabolite (100% for the highest accumulation value) within years or genotypes.

**Figure 7.**
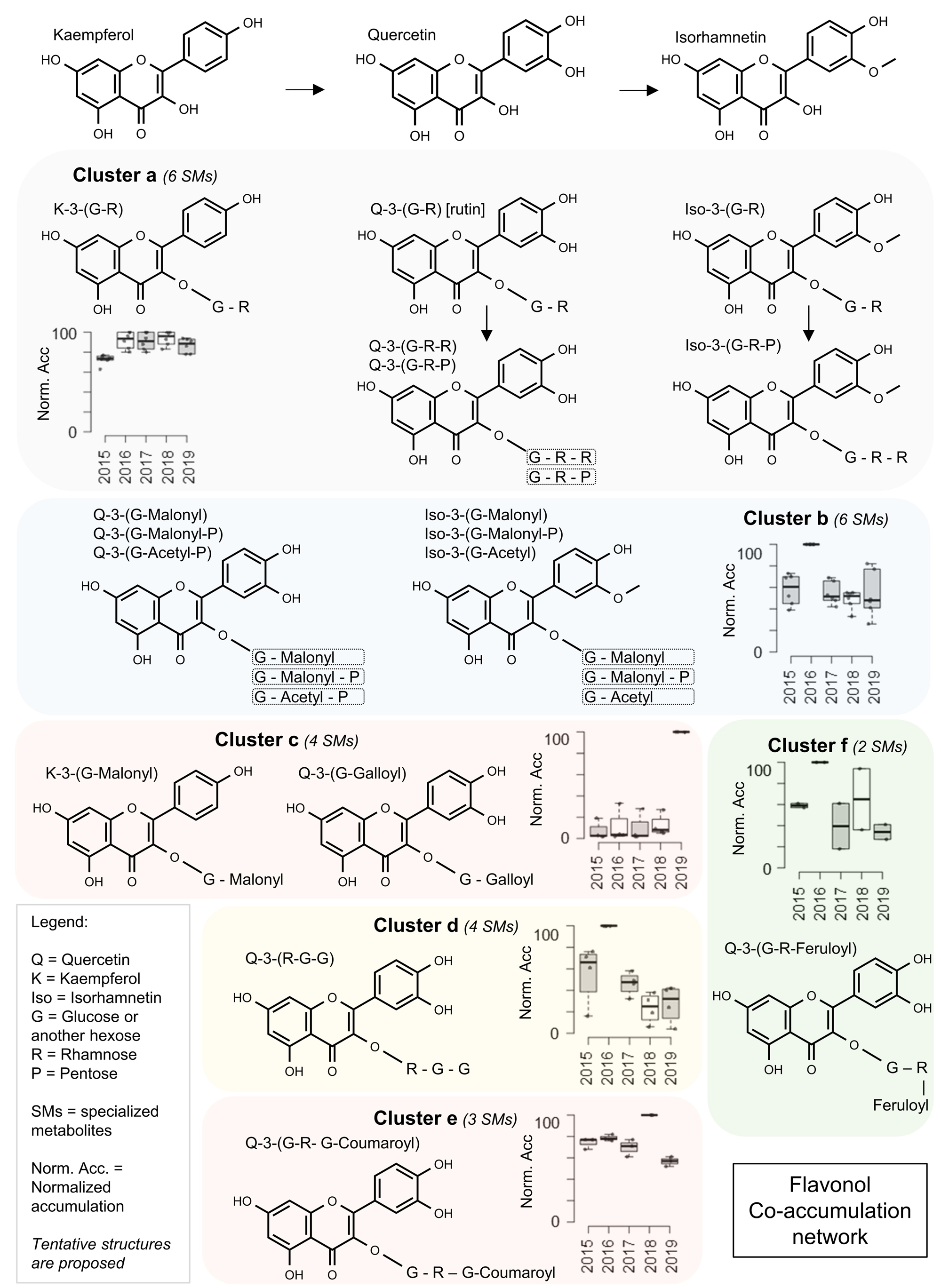
Flavonol-related pathways and clusters Biological pathway representing flavonol-related metabolites. The boxplots represent the accumulation patterns for each metabolite in all years. The accumulation values (boxplots) were normalized and calculated as a percentage related to the sample showing the highest accumulation value for each metabolite (100% for the highest accumulation value) within years or genotypes.

The other annotated enriched metabolic categories, cinnamic acid and glucosinolates, were merged into only one cluster, showing higher accumulation in 2016 (Figure S3). Twenty-nine cinnamic-acid derivatives were incorporated in the network and clustered together (Figure S3a). Most of the cinnamic acids shown in the network derived from sinapic acids and/or sinapine, including different glycosylated forms (Figure S5). Finally, annotated glucosinolates were 9-methylthiononyl glucosinolate, 8-methylsulfinyloctyl glucosinolate and its chain-elongated homologues glucoarabin and glucocamelinin (Figure S6).

Both p_2_a and p_2_c enriched unknown (not annotated) metabolic categories, found using molecular network analyses, were merged into one co-accumulated clusters. As already described for the top-10 most accumulated metabolites (Figure S1), the putative structures of the metabolites belonging to the enriched metabolic categories with no annotation were proposed using SIRIUS 4.0 (Dührkop *et al*., 2019). The metabolites of p_2_a and p_2_c categories were putatively annotated as aminoacids and phenolic acids, respectively. These enriched metabolic categories were characterized by higher accumulation in 2015 and 2016, especially for p_2_c category (Figure S7).

### Comparison of primary metabolites, oil composition and specialized metabolite plasticity in camelina seeds

To gain deeper knowledge on the metabolic composition and the plasticity of seed specialized (e.g. flavan-3-ols, flavonols and glucosinolates) and primary (free aminoacids, sugars, carboxylic acids, fatty acids, lipids) metabolites, additional analyses were performed on selected genotypes. The analysis of primary metabolites (92 metabolites, Table S3), and fatty acids (17 metabolites, Table S4) as well as lipids (1476 metabolic features [Mf], Table S5) were obtained by GC-MS and LC-MS/MS, respectively. Metabolite analyses were carried out on seeds of OMEGA and 789-02 genotypes harvested from 2015 to 2019. These genotypes were chosen based on their contrasted profiles for SMs (Figure 3b, 4b), OMEGA and 789-02 being the ones showing the highest and lowest SM accumulation, respectively. A co-accumulation network analysis (similarly to what has been done previously for SMs; Figure 8a) was performed to calculate the phenotypic plasticity, using a widely used index (Relative Distance Plasticity Index, RDPI; Valladares *et al*., 2006), for the main primary and specialized metabolic categories (Figure 8b; Table S6). The RDPI index allows the quantification of phenotypic plasticity based on phenotypic distances between different individuals of a given species (e.g. camelina plants) exposed to different environments (e.g. 2015 to 2019). While this index has been used to compare plasticity among different species, we applied the method to compare the plasticity of different phenotypes (i.e. the metabolic classes) from the same species and tissue. Co-accumulated (i.e. highly correlated: *r* > 0.8) metabolites were visualised in heatmaps build for each cluster and obtained from the metabolite correlation network (Figure 8a), and the phenotypic plasticity was calculated for each cluster and category (Figure 8b), using the average metabolite intensity (see method for further information). The correlation cut-off has been increased compared to the previous correlation network because of the lower number of genotypes considered and, therefore, of samples used to calculate the correlation.

**Figure 8.**
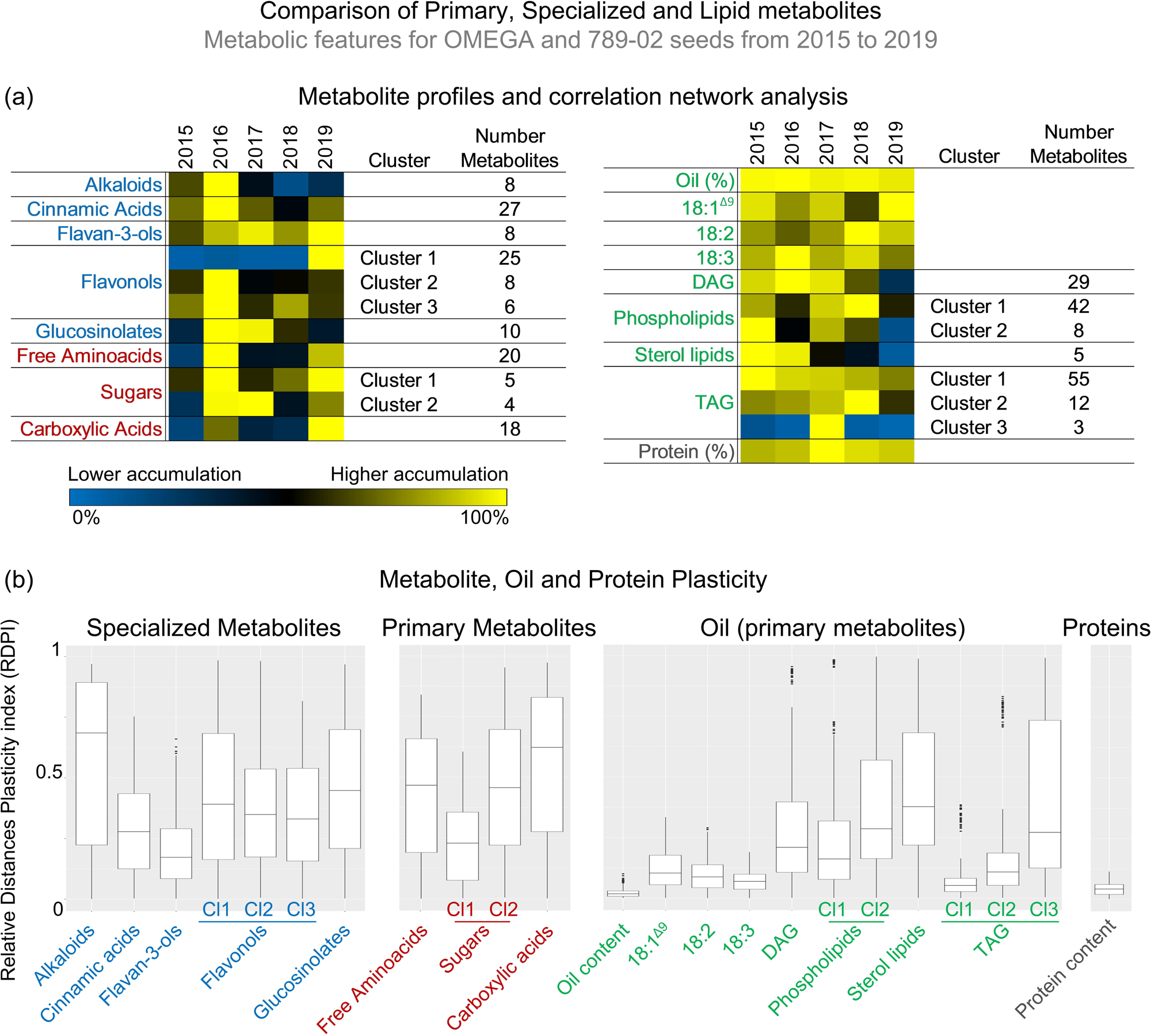
Comparison and phenotypic plasticity of seed primary and specialized metabolites, and oil composition in OMEGA and 789-02 varieties. (a) Heat maps showing the average metabolite accumulation, from 2015 to 2019, for differentially accumulated metabolites belonging to specialized and primary metabolic categories. The accumulation patterns were used to calculate correlations (r > 0.8) among metabolites and assign a metabolic class to a specific cluster. The accumulation values were calculated as a percentage related to the sample showing the highest accumulation value for each metabolite (100% and 0% for yellow and blue colours, respectively) within years or genotypes. (b) Boxplots showing the Relative Distance Plasticity Index (RDPI), representing the phenotypic plasticity for the main specialized and primary (sugars, free aminoacids, carboxylic acids, lipids and proteins) metabolic categories. RDPI index is comprised between 0 (low/no plasticity) and 1 (very high plasticity).

Alkaloids, flavonols, and glucosinolates SMs, together with free aminoacids and carboxylic acids primary metabolites, were the categories showing the largest plasticity level. Conversely, seed oil and protein contents, major fatty acids (18:1^Δ9^, 18:2 and 18:3), most lipid categories (Diacylglycerols, DAG; Triacylglycerols, TAG; Phospholipids) and protein content were characterized by markedly lower plasticity than that just reported for SMs. These categories showed indeed a quite similar accumulation behaviour and no clear metabolic signature across the five years considered in this study. Lower plasticity was also observed for cinnamic acids, flavan-3-ols, and one sugar cluster (which included sucrose, the most abundant sugar). According to metabolite accumulation profiles, 2016, 2017 and 2019 were the years inducing the highest number of specialized and primary metabolites (Figure 8a), including alkaloids (2016), flavonols (2016 and/or 2019), glucosinolates (2016 and 2017), free aminoacids (2016 and 2019), sugars (2016 and, 2017 or 2019), carboxylic acids (2019).

Last, carbon and nitrogen content in seeds of the two genotypes was also analysed. The results for C/N did not reveal a significant impact of genotype nor environment in camelina seeds (Figure S8).

### The environmental regulation of oil composition in camelina seeds

Untargeted lipidomic analyses allowed the annotation of 199 metabolic features (out of 1476), mainly classified into 7 metabolic categories: TAG (85 Mf), Phospholipids (43 Mf), DAG (33 Mf), Fatty acid methyl esters (5 Mf), Sterol lipids (5 Mf), Monoradylglycerols (3 Mf) and Glycerolipids (1 Mf) (Table S5). We also detected (+)-.gamma.-Tocopherol, and minor classes corresponding to Phosphatidylmethanols (8 Mf) and Fatty acid methyl esters, respectively (5 Mf). These last two metabolic classes could be artefacts derived from the extraction protocol that was used. Statistical analyses reveal that 596 (40 %) lipid DAMf were affected by both genotype and environment, belonging to the DAG, TAG, phospholipids and sterol lipids. The heatmaps constructed using the accumulation of metabolites for each category or separately for all metabolites showed that the environmental conditions of 2019 induced lower levels of metabolite accumulation compared to the other years (Figure 9b, Figure S9). In addition, 2017 and 2018 were characterized also by moderately lower level of many DAG and sterol lipids. As already shown by the analysis of phenotypic plasticity, fatty acids (FAs) profiles were stable among the different years, despite the significant but moderate impact of the environment on the accumulation of some major FAs was moderately but significantly affected by the environment (C18:1^Δ9^, C18:2, C18:3) (Figure 9a).

**Figure 9.**
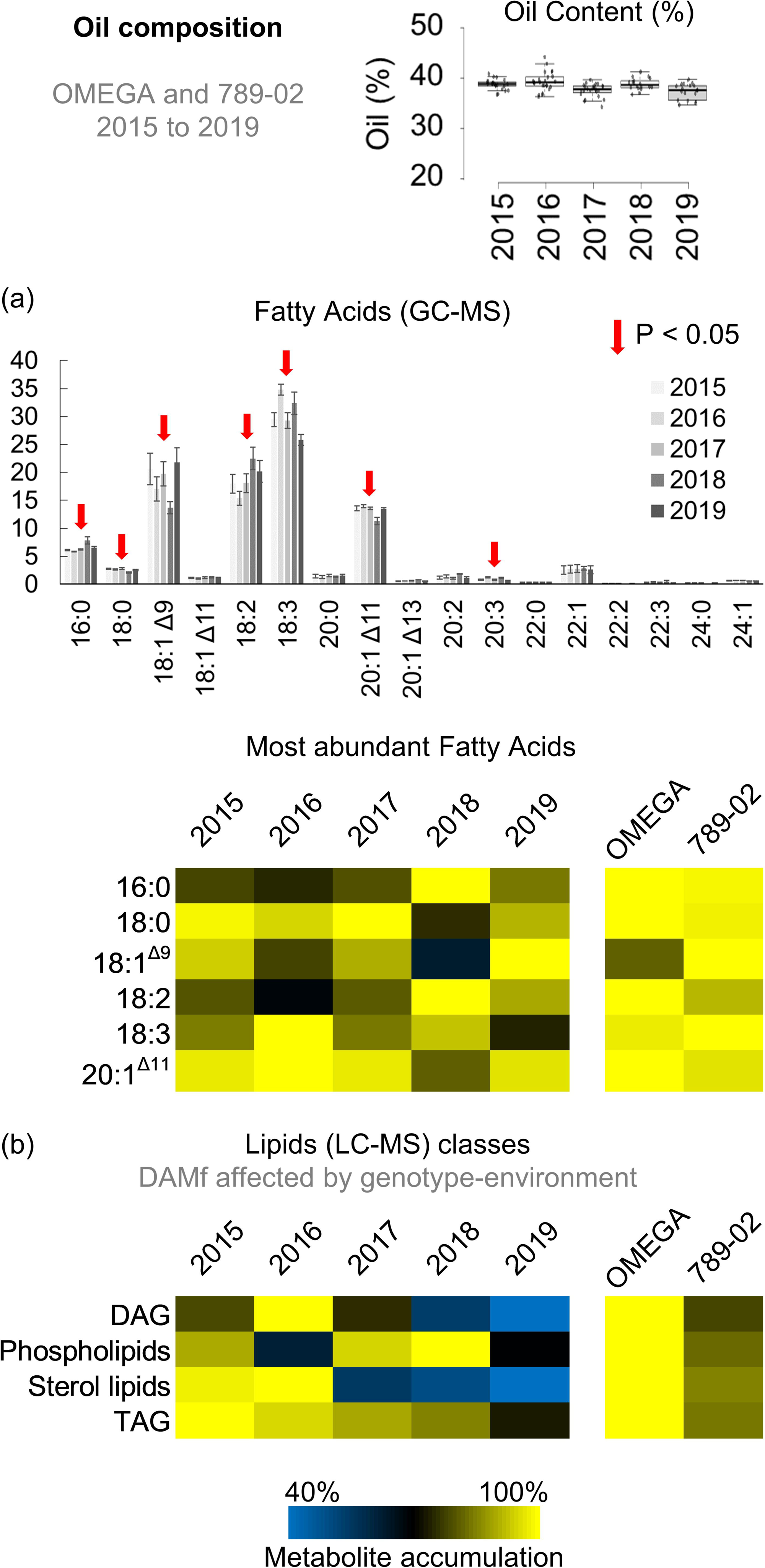
Seed oil composition in OMEGA and 789-02 varieties. (a, b) Fatty acid (a) and lipid (b) profiles and accumulation patterns are shown. Heat maps showing the average metabolite accumulation according to the different years are shown. The accumulation values were calculated as a percentage related to the sample showing the highest accumulation value for each metabolite (100% and 40% for yellow and blue colours, respectively) within years or genotypes.

## DISCUSSION

Phenotypic plasticity refers to the ability of an individual (or a genotype) to cope with environmental changes by modulating one or multiple phenotypic traits (Arnold *et al*., 2019; Fusco and Minelli, 2010). The genetic and environmental impacts on the regulation of gene expression or physiological traits were extensively studied in several plant species and specific tissues or organs (Gruber *et al*., 2013; Dal Santo *et al*., 2016; Dal Santo *et al*., 2013; Janková Drdová *et al*., 2019; Jong *et al*., 2019; Ibañez *et al*., 2017). Conversely, little if any information is available on the overall impact of the environment on the induction or repression of the specialized metabolome, especially in seeds. Studying these processes is particularly interesting in crops, including *Brassicaceae* species such as camelina, a cultivated oilseed crop, whose seeds constitute a valuable feedstock for food, feed and biobased industrial applications. Although it is well known that camelina seeds contain numerous SM classes, most available studies focused on a limited number of compounds, mainly fatty acids or anti-nutritional compounds, such as glucosinolates (Amyot *et al*., 2019).

In the present study, using a molecular network approach (Olivon *et al*., 2018; Elie *et al*., 2019) coupled with manual confirmation of metabolite annotation using molecular structure identification (Dührkop *et al*., 2019), we found a large diversity of SM classes (16 in total) in camelina seeds (Figure 2). The present results highlighted that only 30% of the identified metabolites are annotated or belong to a cluster with annotated metabolites. This is not totally unexpected, since the metabolites present in public databases used for annotation represent only a small portion of the plant metabolome (7-10%) (Chaleckis *et al*., 2019; Szymański *et al*., 2020). It is worth mentioning that the percentage of annotated molecules increased significantly when only the highly accumulated metabolites are considered (Table S1). Despite the large fraction of non-annotate metabolites, using Sirius and molecular network analyses allowed us to increase the percentage of metabolites that were assigned to a metabolic class from 12% (public databases) to 30%. In addition, we significantly expanded the identification of metabolic classes compared with the published untargeted metabolomic analyses on camelina, which mostly focused their attention on annotated metabolites such as sinapine, a few flavonols and flavan-3-ols, and/or glucosinolates (Quéro *et al*., 2016; Berhow *et al*., 2014). Major challenges in the selection of metabolite datasets and their annotation include the duplication of the signal for metabolic features and in-source fragments causing SM miss annotation (Xu *et al*., 2015; Domingo-Almenara *et al*., 2018; Godzien *et al*., 2015). Thus, the impact of in-source ion fragmentation was limited by filtering the metabolic features showing high correlation (r > 0.95) and the same retention time.

Given the five years studied characterized by high meteorological variability and the six genotypes, the trends observed allowed to hypothesize on the major factors impacting the specialized metabolome landscape in camelina seeds. Although the selected genotypes were obtained from different breeding programs and showed major differences in morphology, seed yield, protein and oil content (Righini *et al*., 2019; Zanetti *et al*., 2017; Krzyżaniak *et al*., 2019), metabolite accumulation and diversity can be mainly linked to different weather conditions occurring in each growing season. Indeed, 2016 was characterized by lower temperatures and strikingly higher amounts of precipitation during the seed filling phase, while 2018 and 2019 showed higher temperatures and lower precipitation during the seed filling period. It is worth remarking that while camelina is able to cope quite well with drought stress (Enjalbert *et al*., 2013), its ability to handle high temperatures (Carmo-Silva and Salvucci, 2012) during the seed filling phase is quite limited, and this is in strong agreement with the present results, which highlighted the lowest metabolome induction under those conditions(Figure 3b; 2018 and 2019 growing seasons). In addition, a hailstorm occurred in 2016 prior to harvest, which might have also affected the accumulation of some classes of seed-specialized metabolites. Many of them being well-known antioxidants, which can also act as natural plant defence to adverse and stressful environmental conditions.

Among the annotated metabolites modulated by environment and genotype, flavonols were the most represented class (Figure 4). The flavonol metabolome was induced in several genotypes and years (i.e. by various environmental conditions), as shown by the high number of co-accumulation clusters (six, Figure 6) identified for this category. Hence, since the number of clusters can be directly related to the diversity of a metabolic class, the present data suggest that flavonols are a very plastic category among the SMs observed in camelina seeds. The large accumulation of flavonols in different years could be linked to the ability of plants to respond to different environmental stresses re-directing some biosynthetic pathways. This diversity could be linked to flavonol structure with two aromatic rings and multiple potential decoration sites (glycosylated, methylated and/or acylated; Figures 6 and 7) compared with cinnamic acids (Figure S5) and glucosinolates (Figure S6). The importance of decorations in metabolite activity and plant-interaction with abiotic and biotic factors has been recently highlighted by Behr *et al*. (2020). This is also the case of N-hydroxy-pipecolic acid, whose glycosylation affects plant immunity responses and plant growth (Cai *et al*., 2021; Mohnike *et al*., 2021).

The accumulation of flavonols in response to abiotic stresses has been extensively documented for vegetative tissues (Corso *et al*., 2020; Nakabayashi *et al*., 2014; Tohge *et al*., 2016; Pourcel *et al*., 2007; Muhlemann *et al*., 2018; Emiliani *et al*., 2013), while very few information are available for seeds (Macgregor *et al*., 2015; Corso *et al*., 2020) The antioxidant and protective activities of some flavonols would explain their higher accumulation in 2018 and 2019, which presented the highest temperatures during the seed filling phase, and in 2017, characterized by very limited amounts of precipitation (Figure 1b; Figure 4).

The present analyses suggest that flavonols with different decorations (hydroxylation, glycosylation, methylation and acylation) are differentially accumulated according to environmental conditions (Alseekh, Perez de Souza, *et al*., 2020; Chen *et al*., 2020), especially in response to fluctuating temperatures and high precipitation. In relation to the effects of the environment, the Q -diglycoside – malonyl or feruloyl, and Q-triglycoside, together with acylated and malonylated Q-glycosides were markedly more accumulated in 2016, characterized by high precipitation rate and low temperatures (mix of the two stresses/conditions). Instead, Q and/or K –monoglycoside and -triglycoside (+ malonyl, galloyl or feruloyl) were significantly more accumulated in 2018 and 2019, characterized by higher temperatures (heat stress). Glycosylation, acylation and methylation play crucial roles in stability, solubility and bioavailability of flavonoids. For instance, de-glycosylation is a well-characterized fine-tuned mechanism regulating flavonoid localization, availability, and biological activity (Wang *et al*., 2019). Characterizing the effects of abiotic (e.g. heat or drought stress) and/or biotic stresses on the accumulation of specific flavonoids and related decorations will be essential for understanding the environmental impact on seed specialized metabolism and seed quality.

It has been reported that camelina seed yield and quality (e.g. protein and oil content, oil composition, dormancy or longevity) are affected by meteorological conditions (Zanetti *et al*., 2017; Krzyżaniak *et al*., 2019). Nonetheless, the present study showed that the environment has a limited impact on major seed oil primary metabolites (lipids and fatty acids) and protein contents (Figures 8 and 9), which were characterized by low phenotypic plasticity. Nevertheless, DAG and sterol lipids showed slightly lower accumulation in 2018 and/or 2019 compared with the other years (Figure 9). The impact of abiotic (Hou *et al*., 2016; Shiva *et al*., 2020) and biotic (Jeon *et al*., 2020; Kazaz *et al*., 2021) stresses on lipid accumulation has been already described for vegetative tissues, but the data on seed specialized metabolome are scarce. In particular, osmotic, drought and high temperature stresses enhance the accumulation of phospholipids, including phosphatidylcholine (PC) and phosphatidylethanolamine (PE). The low variability of seed lipids and fatty acid content and composition might be related to camelina domestication history. This species, together with other domesticated *Brassicaceae* (e.g. *Brassica napus*), has been indeed selected for oil production, and showed a higher and less variable seed oil content compared with wild *Brassicaceae* (Barker *et al*., 2007; Sharafi *et al*., 2015).

Conversely, in addition to many SM classes (alkaloids, flavonols, or glucosinolates), a surprisingly high plasticity was observed for free aminoacids, carboxylic acids and some sugars (primary metabolites) in camelina seeds. This suggests that, in addition to the seed specialized metabolome, some metabolic classes belonging to the seed free primary metabolome can be shaped by the environment (Maeda, 2019). This could be due to the fact that many primary metabolites (e.g. free aminoacids) are precursors of major SMs, as in the case of L-phenylalanine for flavonoids (Maeda, 2019; Maeda and Fernie, 2021). A positive induction on the accumulation of seed free primary metabolites has been also observed in Arabidopsis plants subjected to ABA-deficiency, which has been linked to a premature metabolism resumption state (Chauffour *et al*., 2019). Besides free primary metabolites, protein content and turnover might also play a major role in triggering the events leading to successful seed germination and seedling establishment (Galland *et al*., 2014), which could explain the higher stability of camelina seed total protein content highlighted in the current work.

Regarding the differences in metabolite accumulation among the genotypes, both OMEGA and 787-08 showed the highest metabolome induction (Figure 3). Inversely, the genotype 789-02 showed the lowest metabolite accumulation level for most of the studied metabolic categories. Conversely, 787-08 has been previously identified as a cultivar with large seeds, providing good and stable oil yields across different environments (Zanetti *et al*., 2017). Moreover, cultivars 789-02 and 887 are characterized by different seed oil composition (i.e. increased ὠ3/ ὠ6 ratio) particularly suitable to different feed, food and industrial applications. The present results suggest that the agronomic performance of the cultivar 787-08 could be related to an improved genetic combination allowing the diversity and abundance of metabolites. The accordance between metabolite composition and the other traits found in genotypes 787-08 and OMEGA may represent a promising strategy for future breeding in camelina.

In this work were combined two different approaches to annotate and cluster specialized metabolites based on their structure and co-accumulation, respectively. A more precise characterization of new metabolites was achieved by linking their chemical structures to their accumulation patterns. In addition, using fatty acid profiles and untargeted lipidomics it was possible to dissect the impact of the environmental conditions on seed oil composition, important characteristics for a seed oil crop as camelina.

To conclude, major and unexplored effects of the environment on the regulation of the seed specialized metabolome were highlighted, suggesting that seeds might not be exclusively storage and “metabolically static” organs, but they show a very plastic specialized metabolism.

## EXPERIMENTAL PROCEDURES

### Genetic material, field experimental protocol and meteorological parameters

Six different genotypes of camelina were grown at the experimental farm of Bologna University (Italy) located in Cadriano (Bologna, Italy, 44°30′N, 11°23′E, 32 m a.s.l.) during five consecutive years (2015-2019). The soil at Cadriano is characterized as clay-loam (Udic Ustochrepts, mesic). The six tested genotypes were: MIDAS (AAFC, Saskatoon, Canada), OMEGA (University of Poznan, Poland), WUR (Wageningen University and Research, The Netherlands), and 787-08, 789-02 and 887 (Smart Earth Camelina, Saskatoon, Canada). These genotypes were chosen to represent the highest diversity among breeding programs and qualitative seed traits, in particular OMEGA and WUR belong to the breeding programs carried out at Poznan (Poland) and Wageningen (The Netherlands) Universities, respectively. MIDAS has been selected by AAFC (Saskatoon, Canada) to be highly tolerant to disease (i.e,. downey mildew) and with stable productive performance. Cultivars 787-08, 887 and 789-08 derive from the breeding program of Smart Earth Camelina (Canada), and they are improved for: increased seed weight, and stable and high seed yield and oil content (787-08); improved fatty acid composition (887), seed weight and improved fatty acid composition (789-02). Weather parameters (daily minimum, T_min_, maximum, T_max_, and precipitation) were measured and recorded by automated weather stations located at the experimental farm. Camelina was sown each year in late winter/early spring accordingly to the specific meteorological conditions. The same agronomic management for all the trials was adopted, consisting in typical tillage system (ploughing + harrowing), a seeding rate of 500 seeds m^-2^, no irrigation, a top-dressing fertilization with N (50 kg N ha^-1^, as urea at bolting stage), with no pest, disease nor weed chemical control. The experimental design was a randomized complete block with 3 or 4 biological replicates depending on the year. The plants were monitored weekly for their phenological phases, which were surveyed according to the BBCH scale for camelina (Martinelli and Galasso, 2011). At full maturity (residual seed moisture <12%), the central portion of each plot (∼6 m^2^) was cut manually at the soil level, and then threshed using a plot combine. Representative seed samples from individual plots were cleaned at the “Ricerca e Analisi Sementi” laboratory (LaRAS) of DISTAL department (Università di Bologna, IT), and then stored at 4 °C in the dark before shipping to IJPB institute (INRAE-Versailles, FR).

### Protein and Oil extraction and analysis

Oil and protein determinations in all samples have been carried out by AAFC (Saskatoon, Canada), by near-infrared spectroscopy (NIRS) as reported by Zanetti *et al*. (2017). Oil and protein contents are reported as a percentage on a whole seed dry matter (zero moisture) basis.

### Extraction of polar/semipolar specialized metabolites, primary metabolites and lipid fraction

Metabolites were extracted from 60 mg of camelina dry mature seeds (105 samples in total). The metabolite extraction protocol was adapted from Corso *et al*. (2018) and Giavalisco *et al*. (2011).

Briefly, 1 mL of MeOH: Methyl-tert-butyl: H2O (1:3:1), conserved at 4°C, and 200 ng of Apigenin (used as internal standard) were added to each sample, which was then homogenized in 2ml tubes using a FastPrep instrument (1 min, 14,000 rpm). The mixtures were then shaken for 30 min at 4°C using a ThermoMixer™ C (Eppendorf), placed in an ice cooled ultrasonication bath for 15 min and centrifuged for 1 min at the maximum speed to remove debris. The extracted samples were then transferred in new tubes containing 650 mL of MeOH: water (1:3), previously placed at −20°C. The mixtures were centrifuged for 1 min at 14,000 rpm. The addition of MeOH: water (1:3) and the centrifugation led to a phase separation, providing an upper organic phase, containing the lipids, a lower aqueous phase, containing the polar and semi-polar metabolites, and a pellet of starch and proteins. The phase containing the polar / semi-polar metabolites and lipids were dried down in a SpeedVac vacuum concentrator (o/n) and resuspended in 200 μL of ULC/MS grade water (Biosolve).

### UPLC-MS/MS analysis of polar/semipolar specialized metabolites and lipid fractions

Untargeted metabolomic data were acquired using a UHPLC system (Ultimate 3000 Thermo) coupled to quadrupole time of flight mass spectrometer (Q-Tof Impact II Bruker Daltonics, Bremen, Germany). For untargeted polar and semipolar metabolomic analysis, a Nucleoshell RP 18 plus reversed-phase column (2 x 100 mm, 2.7 μm; Macherey-Nagel) was used for chromatographic separation. The mobile phases used for the chromatographic separation were (A) 0.1% formic acid in H_2_O and (B) 0.1% formic acid in acetonitrile. The flow rate was of 400 μL/min and the following gradient was used: 95% of A for 1-min, followed by a linear gradient from 95% A to 80% A from 1 to 3-min, then a linear gradient from 80% A to 75% A from 3 to 8-min, a linear gradient from 75% A to 40% A from 8 to 20-min. 0% of A was hold until 24-min, followed by a linear gradient from 0% A to 95% A from 24 to 27-min. Finally, the column was washed by 30% A at for 3.5 min then re-equilibrated for 3.5-min (35-min total run time).

For untargeted lipidomic analysis, an EC 100/2 Nucleoshell Phenyl-Hexyl column (2 x 100 mm, 2.7 μm; Macherey-Nagel) was used for chromatographic separation. The mobile phases used for the chromatographic separation were (A) H_2_0 + 1% ammonium formate in H_2_O + 0.1% formic acid and (B) Acetonitrile : isopropanol (7 : 3) + 1% of 10mM ammonium formate in H_2_O + 0.1% formic acid. The flow rate was of 400 μL/min and the following gradient was used: 45% of A for 1-min, followed by a linear gradient from 45% A to 30% A from 1 to 2-min, then a linear gradient from 30% A to 15% A from 2 to 7-min, a linear gradient from 15% A to 10% A from 7 to 15-min, a linear gradient from 10% A to 6% A from 15 to 19-min, a linear gradient from 6% A to 2% A from 19 to 26-min. 0% of A was hold until 27-min, followed by a linear gradient from 0% A to 45% A from 27 to 35-min (35-min total run time).

For both untargeted metabolomics and lipidomics, data-dependent acquisition (DDA) methods were used for mass spectrometer data in positive and negative ESI modes using the following parameters: capillary voltage, 4.5kV; nebuliser gas flow, 2.1 bar; dry gas flow, 6 L/min; drying gas in the heated electrospray source temperature, 140°C. Samples were analysed at 8Hz with a mass range of 100 to 1500 *m/z*. Stepping acquisition parameters were created to improve the fragmentation profile with a collision RF from 200 to 700 Vpp, a transfer time from 20 to 70 µs and collision energy from 20 to 40 eV. Each cycle included a MS fullscan and 5 MS/MS CID on the 5 main ions of the previous MS spectrum.

### UPLC-MS/MS data processing

The .d data files (Bruker Daltonics, Bremen, Germany) were converted to .mzXML format using the MSConvert software (ProteoWizard package 3.0, Chambers *et al*., 2012). mzXML data processing, mass detection, chromatogram building, deconvolution, samples alignment, and data export, were performed using MZmine 2.52 software (http://mzmine.github.io/) for both positive and negative data files. The ADAP chromatogram builder (Myers *et al*., 2017) method was used with a minimum group size of scan 3, a group intensity threshold of 1000, a minimum highest intensity of 1500 and *m/z* tolerance of 10 ppm. Deconvolution was performed with the ADAP wavelets algorithm using the following setting: S/N threshold 8, peak duration range = 0.01-1 min RT wavelet range 0.02-0.2 min, MS^2^ scan were paired using a *m/z* tolerance range of 0.01 Da and RT tolerance of 0.4 min. Then, isotopic peak grouper algorithm was used with a *m/z* tolerance of 10 ppm and RT tolerance of 0.4. All the peaks were filtered using feature list row filter keeping only peaks with MS2 scan. The alignment of samples was performed using the join aligner with an *m/z* tolerance of 10 ppm, a weight for *m/z* and RT at 1, a retention time tolerance of 0.3 min. Metabolites accumulation was normalized according to the internal standard (Apigenin) and weight of seeds used for the extraction.

A first research in library with Mzmine was done with identification module and “custom database search” to begin the annotation with our library, currently containing 93 annotations (RT and *m/z*) in positive mode and 61 in negative mode, with RT tolerance of 0.3 min and *m/z* tolerance of 0.0025 Da or 6 ppm.

### Molecular networking of untargeted metabolomic data

Molecular networks were generated with MetGem software (Olivon *et al*., 2018; https://metgem.github.io) using the .mgf and .csv files obtained with MZmine2 analysis. The molecular network was optimized for the ESI+ and ESI-datasets and different cosine similarity score thresholds were tested. ESI- and ESI+ molecular networks were generated using cosine score thresholds of 0.75 and 0.8, respectively. Molecular networks were exported to Cytoscape software (Shannon *et al*., 2003; https://cytoscape.org/) to format the metabolic categories.

### Metabolite annotation of untargeted metabolomic data

Metabolite annotation was performed in four consecutive steps. First, the obtained RT and *m/z* data of each feature were compared with our home-made library containing more than 150 standards or experimental common features (RT, *m/z*). Second, the ESI- and ESI+ metabolomic data used for molecular network analyses were searched against the available MS^2^ spectral libraries (Massbank NA, GNPS Public Spectral Library, NIST14 Tandem, NIH Natural Product and MS-Dial), with absolute *m/z* tolerance of 0.02, 4 minimum matched peaks and minimal cosine score of 0.8. Third, not-annotated metabolites that belong to molecular network clusters containing annotated metabolites from step 1 and 2 were assigned to the same chemical family. Finally, for metabolites that had no or unclear annotation, Sirius software (https://bio.informatik.uni-jena.de/software/sirius/) was used. Sirius is based on machine learning techniques that used available chemical structures and MS/MS data from chemical databanks to propose structures of unknown compounds. Raw data were normalised on the internal standard (Apigenin) and weight of seeds used for the extraction.

### Metabolite annotation of untargeted lipidomic data

Lipid metabolites annotation was performed in different consecutive steps. First, the ESI- and ESI+ metabolomic data used for molecular network analyses were searched against the available MS^2^ spectral libraries (Massbank NA, GNPS Public Spectral Library, NIST14 Tandem and NIH Natural Product), with absolute *m/z* tolerance of 0.02; 4 minimum matched peaks and minimal cosine score of 0.65. Second, in the different clusters of the molecular network, the result of the database search was validated using the different specific fragments and neutral loss for the different lipid class with their MS^2^ spectrum (*<u>Lipid Class Specific Fragments - Lipidomics-Standards-Initiative (LSI)</u>*). If the database search result was valid, annotation of other features was performed by stepwise comparison from the valid lipid metabolite. Finally, for the cluster of molecular network that had no data base search result, Sirius software (https://bio.informatik.uni-jena.de/software/sirius/) was used.

### Chemical structures drawing

Specialized metabolites structures were drawn using ChemDraw 19.0 with default parameters

### Fatty Acids analysis

Fatty acid composition of the seed oil was determined using the same protocol described in Zanetti *et al*. (2017)

### Primary Metabolites: GC-MS data analysis and processing

The lower aqueous phase that was obtained after phase separation of the extracted samples was used to analyse primary metabolites via GC-MS.

GC-MS analyses and data processing were performed according to Fiehn (2006), Fiehn *et al*. (2008) and Ponnaiah *et al*. (2019). Briefly, 150µl of the extraction solution were taken and dry in a Speed-Vac evaporator for 2 h at 30°C before adding 10 µL of 20 mg.mL-1 methoxyamine in pyridine to the samples. The reaction was performed for 90 min at 30°C under continuous shaking in an Eppendorf thermomixer. 90 µL N-methyl-N-trimethylsilyl-trifluoroacetamide (MSTFA) (Regis Technologies, Morton Grove, IL, USA) were then added and the reaction continued for 30 min at 37°C. After cooling, all the samples were transferred to an Agilent vial for injection. At 4 h after derivatization, 1 µL of sample was injected in splitless mode on an Agilent 7890B gas chromatograph coupled to an Agilent 5977A mass spectrometer. The column was an Rxi-5SilMS from Restek (30 m with 10 m Integra-Guard column). An injection in split mode with a ratio of 1:30 was systematically performed for saturated compounds quantification. Oven temperature ramp was 60°C for 1 min then 10°C min-1 to 325°C for 10 min. Helium constant flow was 1.1 mL.min-1. Temperatures were the following: injector: 250°C, transfer line: 290°C, source: 230°C and quadrupole 150°C. The quadrupole mass spectrometer was switched on after a 5.90 min solvent delay time, scanning from 50 to 600 *m/z*. Absolute retention times were locked to the internal standard d27-myristic acid using the RTL system provided in Agilent’s Masshunter software. Retention time locking reduces run-to-run retention time variation. Samples were randomized. A fatty acid methyl esters mix (C8, C9, C10, C12, C14, C16, C18, C20, C22, C24, C26, C28, C30) was injected at the beginning of analysis for external RI calibration. The Agilent Fiehn GC/MS Metabolomics RTL Library (version June 2008) was employed for metabolite identifications. Peak areas determined with the Masshunter Quantitative Analysis (Agilent Technologies, Santa Clara, CA, USA) in splitless and split 30 modes. Resulting areas were compiled into one single MS Excel file for comparison. Peak areas were normalized to Ribitol and Dry Weight. A total of 92 unique metabolites were identified are expressed in arbitrary units (semi-quantitative determination).

### Statistical and enrichment analyses on polar and semi/polar specialized metabolomic data

Normalised metabolite accumulations were expressed as Relative Units (RU). Only metabolites having values higher than 0.05 RU in at least 1/6 of the samples (corresponding to the number of genotypes) were considered. For these 1481 metabolic features, accumulation was modeled with a linear model to decompose the averaged accumulation by a genotype effect plus a year effect and an interaction between the genotype and the year. Several questions were addressed with this model (i) the differences of accumulation between two genotypes averaged on years or given a year (ii) the differences of accumulation between two years averaged on genotypes or given a genotype (iii) the interactions between genotypes and years by testing if the difference of accumulation between two years in a given genotype is the same for a second genotype.

All these questions were written as contrasts in the linear model. P-values were adjusted across the metabolites for each question to control the false discovery rate (FDR), and a difference of accumulation for a metabolite was declared significant if its adjusted p-value was lower than 0.05. A Venn diagram, with the number of DAM depending on the year, genotype and genotype X year interaction, was then made (http://jvenn.toulouse.inra.fr/app/example.html).

Enrichment analysis was done by using a hypergeometric test and the FDR was controlled across all the terms at level 0.05. The metabolite reference set was defined from the annotated and unknown metabolic categories (Table S1, column ‘Metabolic category’). The analyses was performed with the software R. The contrasts and the enrichment tests were written according to the functions of the tool DiCoExpress (Lambert *et al*., 2020).

### Statistical analyses on lipids and primary metabolite data

Similarly to polar/semipolar specialized metabolites, lipids and primary metabolites data were normalized to an internal standard. Differentially Accumulated Metabolites (DAM) or metabolic features (DAMf) were then identified following the same statistical pipeline used for polar/semipolar metabolites. DAM and/or DAMf affected by the environment and/or the genotype were then used for further computational analyses and data interpretation.

### Total carbon and nitrogen contents

Three milligram of seeds were placed in tin capsules and burned in an elemental analyser (Pyrocube, Elementar, Lyon, France). Separation of resulting N_2_ and CO_2_ gas was performed on two gas-selective columns with Thermal Conductivity Detector (TCD). They can be quantified against standards (ammonium sulfate 21.2% N, benzoic acid 68.85 % C, glutamic acid 9.51% N / 40.82% C and glutamine 19.17% N / 41.09% C). The elemental carbon or nitrogen content was given in % (mass fraction).

### Heatmap, hierarchical clustering, metabolites co-accumulation analysis and calculation of phenotypic plasticity

The heatmaps and hierarchical clustering analyses of metabolomic data were carried out using the ‘heatmap.2’ function of GPLOTS R package (https://cran.r-project.org/bin/windows/base/). The metabolite value used for the heatmaps were calculated as percentage related to the sample showing the highest accumulation value for each metabolite (maximum and minimum accumulation for yellow and blue colours, respectively; specific values are reporter for each heatmap).

Correlation was calculated among metabolites from enriched metabolic classes or with the average accumulation patterns of all annotated metabolic classes using ‘ExpressionCorrelation’ app of Cytoscape software (Shannon *et al*., 2003). All 105 samples present in the dataset were used to calculate correlations among metabolites. Metabolite co-accumulation networks were visualised using Cytoscape software.

To measure phenotypic plasticity, the Relative Distance Plasticity Index (RDPI; Valladares *et al*., 2006) was calculated for specialized and primary metabolic categories using the *Plasticity* R package (https://github.com/ameztegui/Plasticity).

### Statistical analyses on physiological and seed quality parameters

To assess statistical differences in physiological parameters (yield), protein and oil content, and fatty acids composition among the different years (2015, 2016, 2017, 2018, 2019) or *Camelina sativa* accessions (MIDAS, OMEGA, WUR, 787.08, 789.02 and 887), an ANOVA with Tukey’s range test (*P*≤0.05) post-hoc correction was performed using the AGRICOLAE R package.

## ACKNOWLEDGEMENTS

This work was supported by the “Département de Biologie et Amélioration des Plantes” (BAP) of INRAE (grant “Appel à projets scientifiques BAP 2020” to M.C.) and by a grant (“Seed MetSpe”) of the Labex Saclay Plant Sciences-SPS (AN-10-LABX-0040-SPS to L.L. and G.M.). IJPB and IPS2 benefit from the support of Labex Saclay Plant Sciences-SPS (ANR-17-EUR-0007). This work has benefited from the support of IJPB’s Plant Observatory technological platforms and IPS2’s metabolomic platform. Authors would like to thank Angela Vecchi for the technical supervision of the field trials in Bologna, and the LaRAS personnel for the technical supervision of the seed sample preparation. Authors thank also the seed providers of all the tested genotypes, and AAFC (Saskatoon, Canada) for running the oil and protein content determinations on camelina seeds. We would like to thank as well Dr. Sébastien Baud, Dr Grégory Mouille and Prof. Loïc Rajjou (IJPB, INRAE) for the constructive discussion on seed fatty acids, primary and specialized metabolites.

## DATA AVAILABILITY

The metabolomic data and metadata, and script used for the statistical analyses have been deposited at the INRAE data repository portal (data INRAE) with the following identifiers:

1. LC-MS/MS untargeted metabolomics, https://doi.org/10.15454/ATTENN.
2. LC-MS/MS untargeted lipidomics, https://doi.org/10.15454/ODKCCS.
3. GC-MS metabolomics, https://doi.org/10.15454/XYNDVB.
4. R script used for the statistical analyses, https://doi.org/10.15454/A3QKZI.

## AUTHOR CONTRIBUTIONS

MC and FZ designed the research. MC, FZ, AM and LL directed the research. FZ and AM coordinated field experiments and furnished the seed samples for the analyses. SB and FP generated the LC-MS/MS data, performed the initial quality analyses on the metabolomic data and the metabolites annotation with SIRIUS. FZ and AM performed the analysis of fatty acids. JCT performed the untargeted lipidomic analyses (LC-MS/MS). CM and BK performed the analyses on primary metabolites (GC-MS). MC, SB and LB performed the molecular network analyses. MC, LB, ED and MLM performed the computational analyses on metabolomic data. MC and LB generated all Fig.s and supplemental material. MC and LB wrote the paper, which was edited by FZ, AM and LL. All the authors commented on and approved the manuscript.

## SUPPORTING INFORMATION

Table S1. Annotation and information on the Camelina sativa specialized metabolome.

Table S2. Relative accumulation of camelina seed specialized metabolites (LC-MS/MS).

Table S3. Diversity and accumulation of seed primary metabolites (GC-MS) for OMEGA and 789-02 camelina genotypes.

Table S4. Diversity and accumulation of seed fatty acids (GC-MS) for OMEGA and 789-02 camelina genotypes.

Table S5. Diversity and accumulation of seed lipids (LC-MS/MS) for OMEGA and 789-02 camelina genotypes.

Table S6. Co-accumulation networks of primary and specialized metabolites for OMEGA and 789-02 camelina genotypes.

Figure S1. Top-10 most accumulated specialized metabolites in camelina seeds.

Figure S2. Induced and repressed metabolites identified in all comparisons for genotype and environment factors.

Figure S3. Cinnamic acid and glucosinolate co-accumulation networks.

Figure S4. Co-accumulation networks constructed with the unknown metabolic categories

Figure S5. Cinnamic acids-related pathways and clusters

Figure S6. Glucosinolate structures and clusters

Figure S7. Cluster accumulation behaviours, putative structures and annotations for the enriched but unknown metabolic classes.

Figure S8. Elemental carbon and nitrogen content in 789-02 and OMEGA seeds.

Figure S9. Heatmap representing lipids influenced by the environment and the genotype.

