## Supplemental Figures for "Untargeted metabolomic analyses reveal the diversity and plasticity of the specialized metabolome in seeds of different *Camelina sativa* genotypes"

**SUPPORTING INFORMATION**

**Table S1.** Annotation and information on the Camelina sativa specialized metabolome.

Identification (ID), ionization mode (ESI), *m/z*, retention time (RT), Molecular Network clusters, metabolic category (based on molecular network clusters and annotation based on standards and according to public databases), and source of the annotation are indicated for each metabolite. n = metabolite or cluster from negative ionization mode, p = metabolite or cluster from positive ionization mode. Legend for the annotation of metabolites: Q = quercetin, K = kampferol, Iso = isorhamnetin, G = glucose or other hexose, R = rhamnose, P = pentose

**Table S2.** Relative accumulation of camelina seed specialized metabolites (LC-MS/MS).

Data are showed for the six genotypes (MIDAS, OMEGA, WUR, 787-08, 789-02, 887), the five years (2015-2019) and biological replicates (1, 2, 3, 4) considered in this study. The CQs were obtained by mixing all the samples considered in the analyses

**Table S3.** Diversity and accumulation of seed primary metabolites (GC-MS) for OMEGA and 789-02 camelina genotypes.

**Table S4.** Diversity and accumulation of seed fatty acids (GC-MS) for OMEGA and 789-02 camelina genotypes.

Fatty acid identification and accumulation data are shown for OMEGA and 789-02 varieties, five years (2015-2019) and three/four biological replicates (1, 2, 3, 4). NA = not available.

**Table S5.** Diversity and accumulation of seed lipids (LC-MS/MS) for OMEGA and 789-02 camelina genotypes.

Identification (ID), ionization mode (ESI), *m/z*, retention time (RT), Molecular Network clusters, metabolic class (based on molecular network clusters) and annotation based on standards or/and according to public databases, are indicated for each metabolite. n = metabolite or cluster from negative ionization mode, p = metabolite or cluster from positive ionization mode. Metabolite accumulation data are shown for OMEGA and 789-02 varieties, five years (2015-2019) and three/four biological replicates (1, 2, 3, 4).

**Table S6.** Co-accumulation networks of primary and specialized metabolites for OMEGA and 789-02 camelina genotypes. A correlation network was constructed using metabolite intensities for the main specialized and primary metabolites for OMEGA and 789-02 camelina genotypes Identification (ID), Metabolic class (based on molecular network clusters) and cluster number are indicated for each metabolite represented in the network ($r > 0.8$).

**Figure S1.** Top-10 most accumulated specialized metabolites in camelina seeds.
(a) Box plot showing the relative accumulation of the top-10 most accumulated specialized metabolites. Centre lines show the medians; box limits indicate the 25th and 75th percentiles as determined by R software; whiskers extend 1.5 times the interquartile range from the 25th and 75th percentiles, outliers are represented by dots; data points are plotted as open circles. n = 30 sample points. The metabolite levels are represented as relative units normalized according to the internal standard (Apigenin) and relative to the seed weight.
(b) Structures of the most accumulated metabolites that were annotated using public databases.

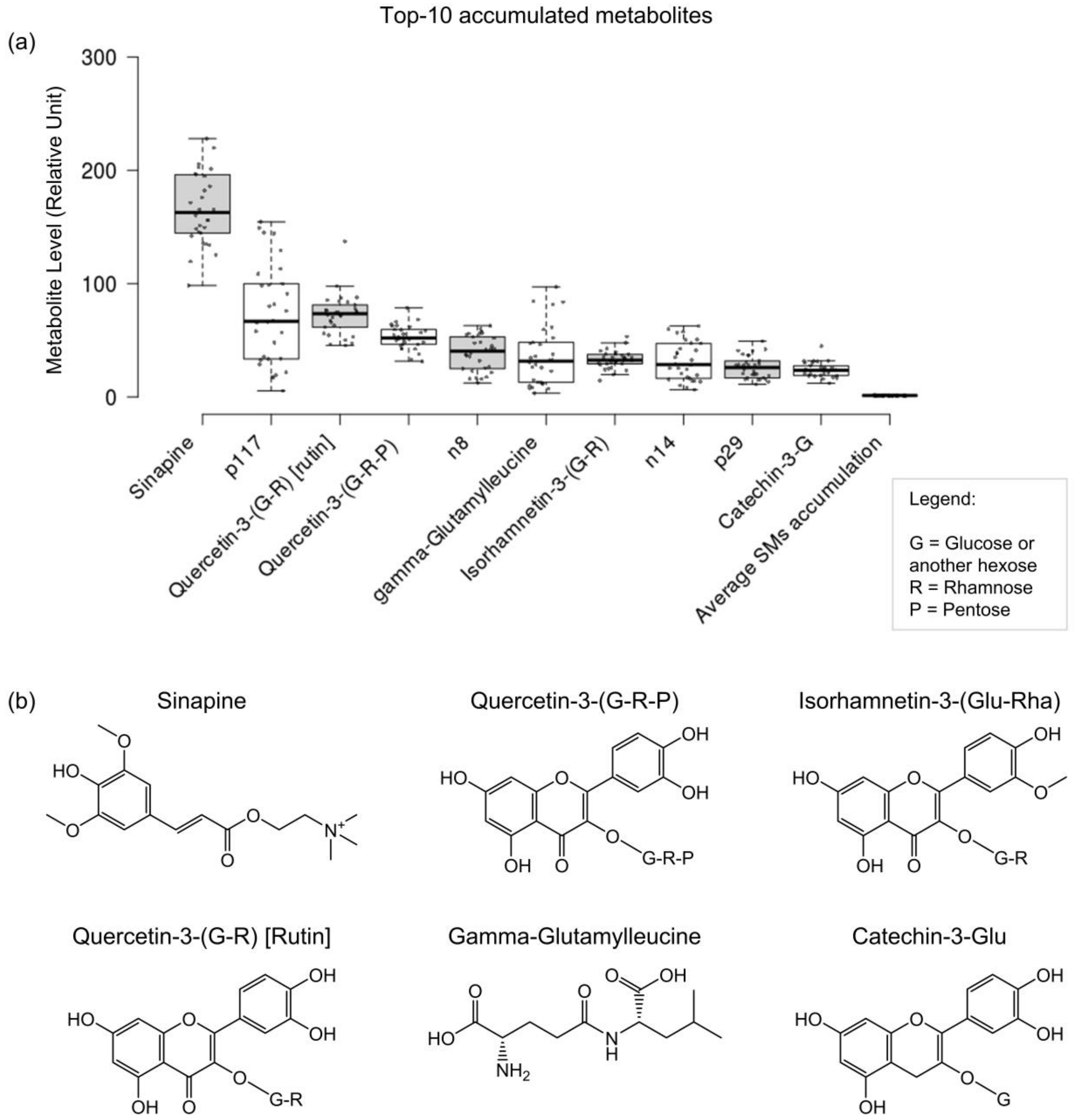

**Figure S2.** Induced and repressed metabolites identified in all comparisons for genotype and environment factors.

Differentially accumulated metabolites in all comparisons for the genotype and the environment factors. Induced and repressed metabolites were calculated for each comparison: e.g. for the 2015 vs 2016 comparison, 111 metabolites were more accumulated (induced) in 2015 than in 2016, and 587 less accumulated (repressed) in 2015 than in 2016.

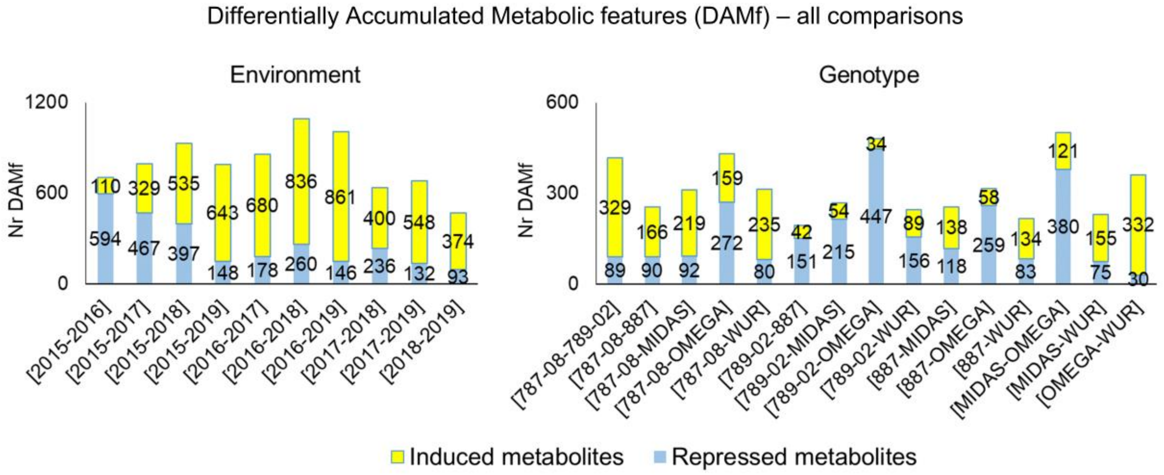

**Figure S3.** Cinnamic acid and glucosinolate co-accumulation networks.
(a, b) Clusters and accumulation patterns are shown for the cinnamic acid (a) and glucosinolate (b) metabolites present in the co-accumulation network. The boxplots represent the accumulation patterns for each metabolite in all years and genotypes. The *m/z*, retention time, MetGem cluster and annotations are also shown for each metabolite.

Metabolite Correlation Networks
Annotated enriched-metabolic categories

(a) **Cinnamic acids**

| ID | mz | RT | Cluster MetGem | Annotation |
|---|---|---|---|---|
| n155 | 663.2 | 4.8 | n_6 | |
| n158 | 385.1 | 3.8 | n_41 | Synapoyl-G |
| n169 | 493.1 | 5.0 | n_6 | |
| n20 | 693.2 | 4.7 | n_6 | |
| n242 | 457.1 | 3.2 | n_1_d | Coumaric acid-ApiosylG |
| n337 | 380.1 | 3.9 | n_1_d | |
| n418 | 665.2 | 5.7 | n_6 | |
| n431 | 723.2 | 5.4 | n_6 | |
| n437 | 591.2 | 5.6 | n_6 | |
| 431 | 519.2 | 3.9 | n_6 | |
| n48 | 431.1 | 1.9 | n_1_d | |
| n630 | 591.2 | 7.1 | n_6 | |
| n695 | 341.1 | 4.1 | n_1_d | |
| n696 | 325.1 | 2.9 | n_1_d | Coumaric acid-G |
| n701 | 367.1 | 4.2 | n_17 | Feruloylquinic acid |
| n790 | 561.2 | 6.6 | n_6 | |
| n880 | 366.1 | 1.1 | n_1_d | |
| p1026 | 307.1 | 3.7 | p_10 | |
| p142 | 296.1 | 2.9 | p_10 | Feruloylcholine |
| p1493 | 446.2 | 3.8 | p_10 | 1-[(3,4-Dimethoxyphenyl)-(3,4,5-trimethoxyphenyl)methyl]pipecolic |
| p1995 | 294.2 | 4.3 | p_10 | |
| p1997 | 250.1 | 3.2 | p_10 | 4-Coumaroylcholine |
| p332 | 476.2 | 3.9 | p_10 | |
| p51 | 280.2 | 3.5 | p_10 | |
| p591 | 224.1 | 1.5 | p_10 | 4-Hydroxybenzoylcholine |
| p593 | 638.3 | 3.3 | p_10 | GAG-ß-feruloyllcholine ether-G |
| p634 | 472.2 | 2.7 | p_10 | Sinapine-4-G |
| p674 | 354.2 | 3.2 | p_10 | 3-(3-Hydroxypropylamino)propyl 3,4,5-trimethoxycinnamate |
| p793 | 390.1 | 4.2 | p_377 | |

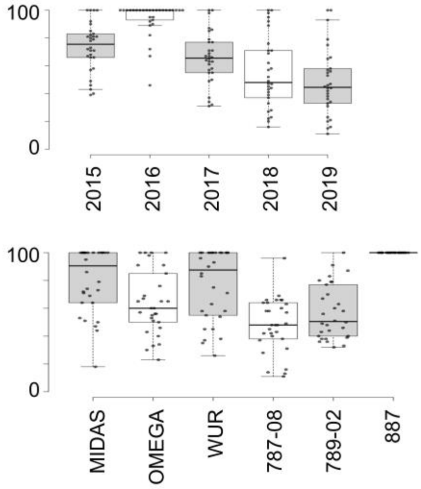


(b) **Glucosinolates**

| ID | mz | RT | Cluster MetGem | Annotation |
|---|---|---|---|---|
| n14 | 506.1 | 4.0 | n_5 | Glucoarabin |
| n258 | 506.1 | 10.0 | n_5 | |
| n283 | 492.1 | 3.5 | n_5 | Methylsulfinyloctyl glucosinolate |
| n381 | 536.1 | 3.9 | n_5 | |
| n398 | 550.1 | 3.8 | n_5 | |
| n405 | 490.1 | 5.1 | n_5 | 9-Methylthiononyl Glucosinolate |
| n420 | 504.1 | 6.4 | n_5 | |
| n511 | 534.2 | 13.7 | n_5 | |
| n590 | 522.1 | 2.6 | n_5 | |
| n8 | 520.1 | 4.6 | n_5 | Glucocamelinin |

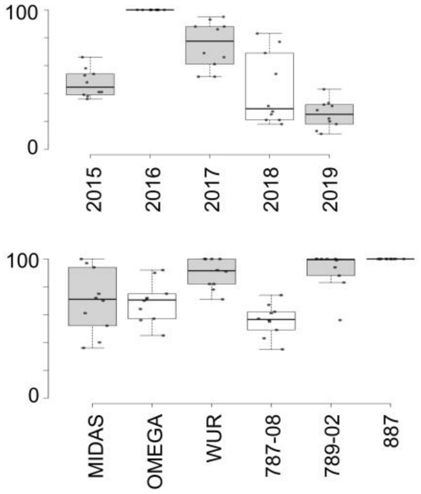


Legend:
G = Glucoside
GAG = Guaiasylglycerol

**Figure S4.** Co-accumulation networks constructed with the unknown metabolic categories (a, b, c) Clusters and accumulation profiles are shown for the metabolites that belong to the p_2_a (a), p_2_c (b) and p_16 (c) unknown categories and present in the co-accumulation networks. The boxplots represent the accumulation patterns for each metabolite in all years and genotypes. The *m/z*, retention time, MetGem cluster and annotations are also shown for each metabolite.

Metabolite Correlation Networks
Unknown enriched-metabolic categories

(a) **p_2_a**

| **ID** | **mz** | **RT** | **Cluster MetGem** |
|---|---|---|---|
| p790 | 323.1 | 3.9 | p_2_a |
| p547 | 324.1 | 1.2 | p_2_a |
| p739 | 324.1 | 1.4 | p_2_a |
| p1078 | 338.1 | 2.3 | p_2_a |
| p613 | 339.1 | 2.9 | p_2_a |
| p617 | 349.1 | 5.1 | p_2_a |
| p536 | 362.1 | 4.2 | p_2_a |
| p1753 | 365.1 | 3.7 | p_2_a |
| p525 | 366.2 | 4.1 | p_2_a |
| p1093 | 369.1 | 3.3 | p_2_a |
| p658 | 391.2 | 10.2 | p_2_a |
| p565 | 392.1 | 5.2 | p_2_a |
| p1784 | 404.2 | 7.5 | p_2_a |
| p2031 | 407.2 | 6.1 | p_2_a |
| p540 | 420.2 | 8.7 | p_2_a |
| p1876 | 431.1 | 4.8 | p_2_a |
| p1543 | 441.1 | 9.8 | p_2_a |
| p570 | 454.1 | 7 | p_2_a |
| p1153 | 583.2 | 6.1 | p_2_a |

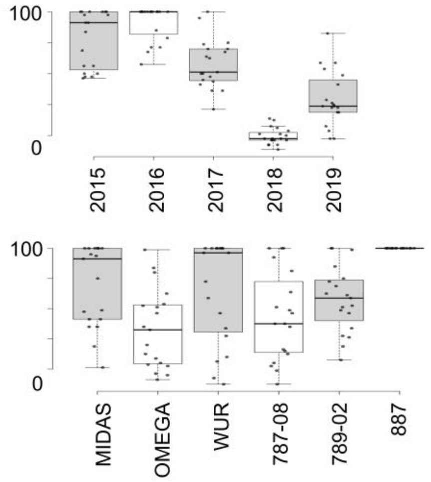


(b) **p_2_c**

| **ID** | **mz** | **RT** | **Cluster MetGem** |
|---|---|---|---|
| p695 | 307.1 | 2.9 | p_2_c |
| p346 | 291.1 | 4.3 | p_2_c |
| p1544 | 642.2 | 6.4 | p_2_c |
| p874 | 307.1 | 4.9 | p_2_c |
| p1562 | 305.1 | 5.5 | p_2_c |
| p1567 | 268.1 | 3.2 | p_2_c |
| p1953 | 441.1 | 6 | p_2_c |
| p1368 | 390.2 | 5.8 | p_2_c |
| p862 | 468.1 | 9.4 | p_2_c |

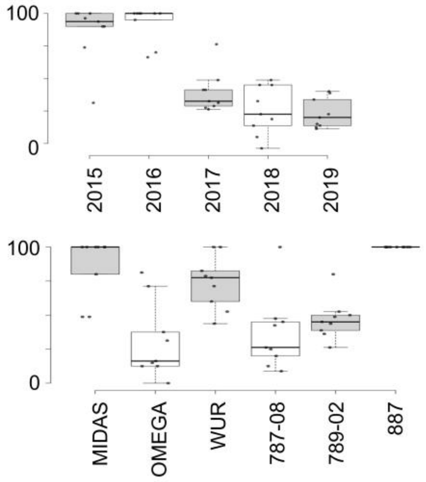

**Figure S5.** Cinnamic acids-related pathways and clusters

Biological pathway representing the cinnamic acid-related metabolites. The boxplots represent the accumulation patterns for each metabolite in all years.

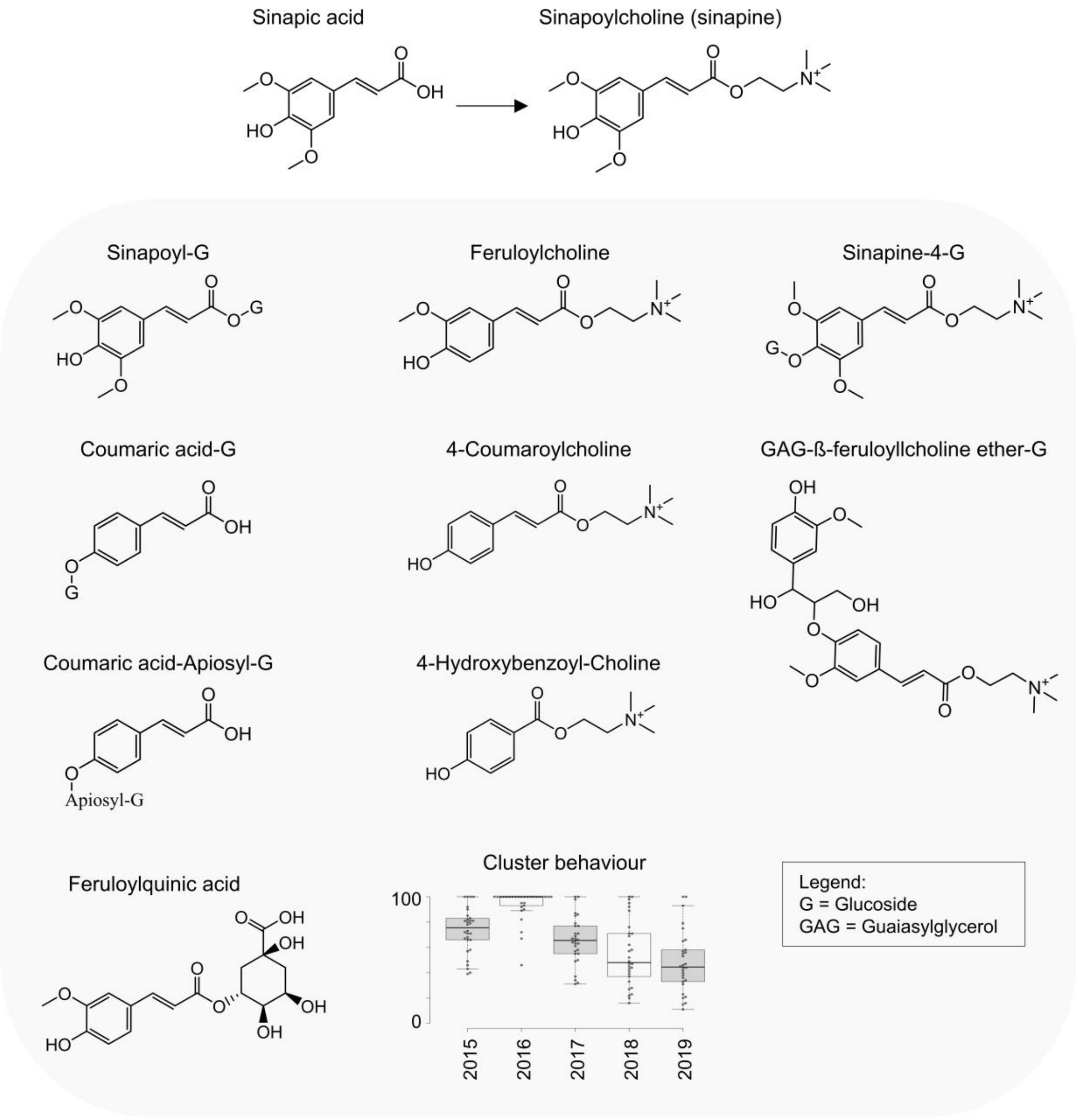

**Figure S6.** Glucosinolate structures and clusters

Chemical structure representing the annotated glucosinolate metabolite. The boxplots represent the accumulation patterns for each metabolite in all years.

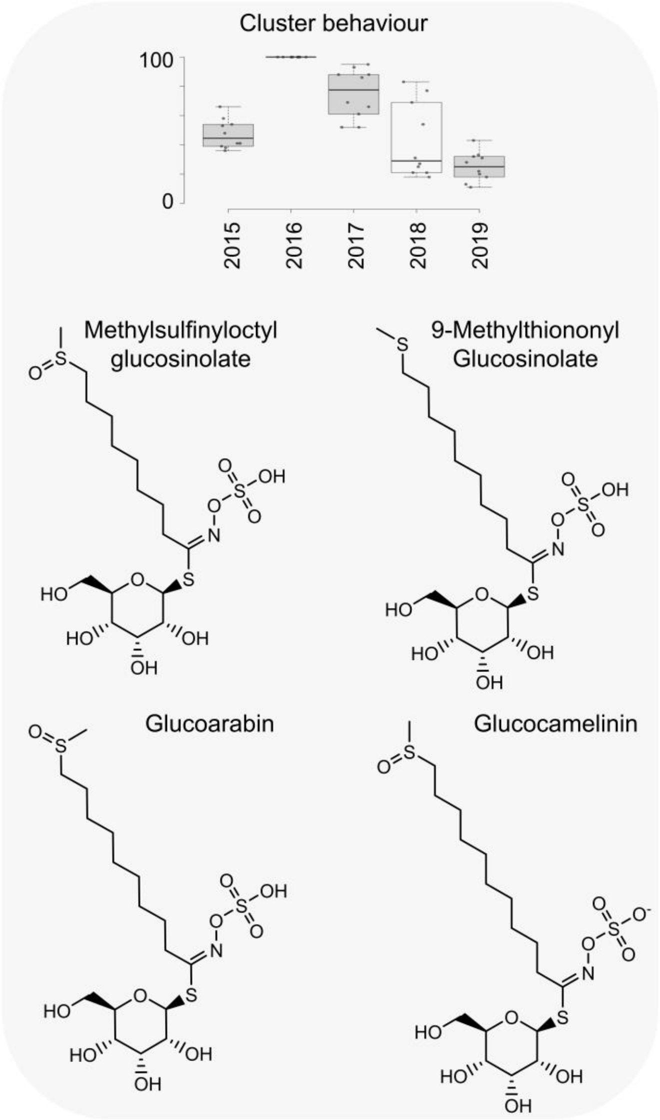

**Figure S7.** Cluster accumulation behaviours, putative structures and annotations for the enriched but unknown metabolic classes.

Putative chemical structures representing the unknown metabolic classes that showed an enrichment. The boxplots represent the accumulation patterns for each metabolite in all years.

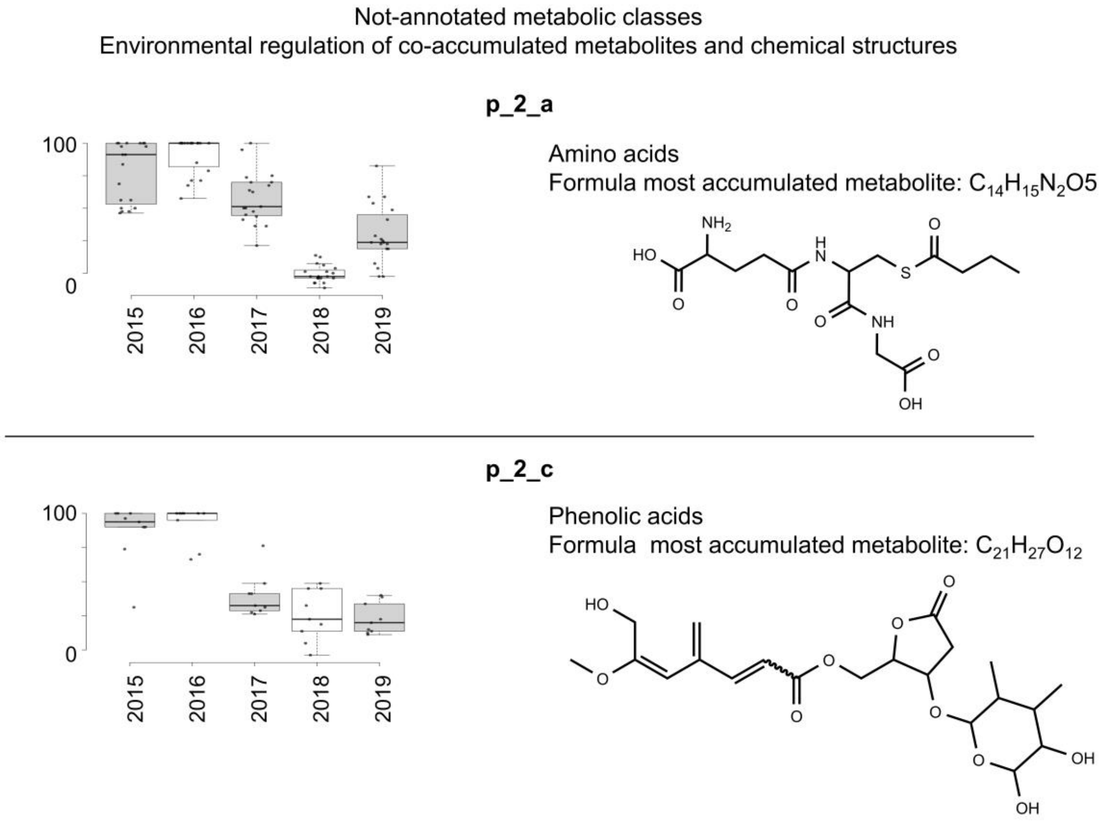

**Figure S8.** Elemental carbon and nitrogen content in 789-02 and OMEGA seeds. C/N ration is shown for each year (2015 to 2019) and genotype considered. ns = not significant (Tukey's range test).

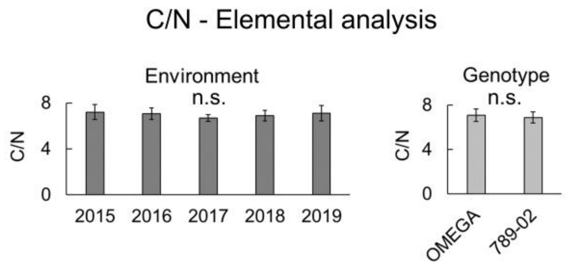

**Figure S9.** Heatmap representing lipids influenced by the environment and the genotype. The accumulation patterns of annotated DAMf affected by both the genotype and the environment are shown. The accumulation values were calculated as a percentage relative to the sample showing the highest accumulation value for each metabolite (100% and 0% for yellow and blue colours, respectively) within years or genotypes.

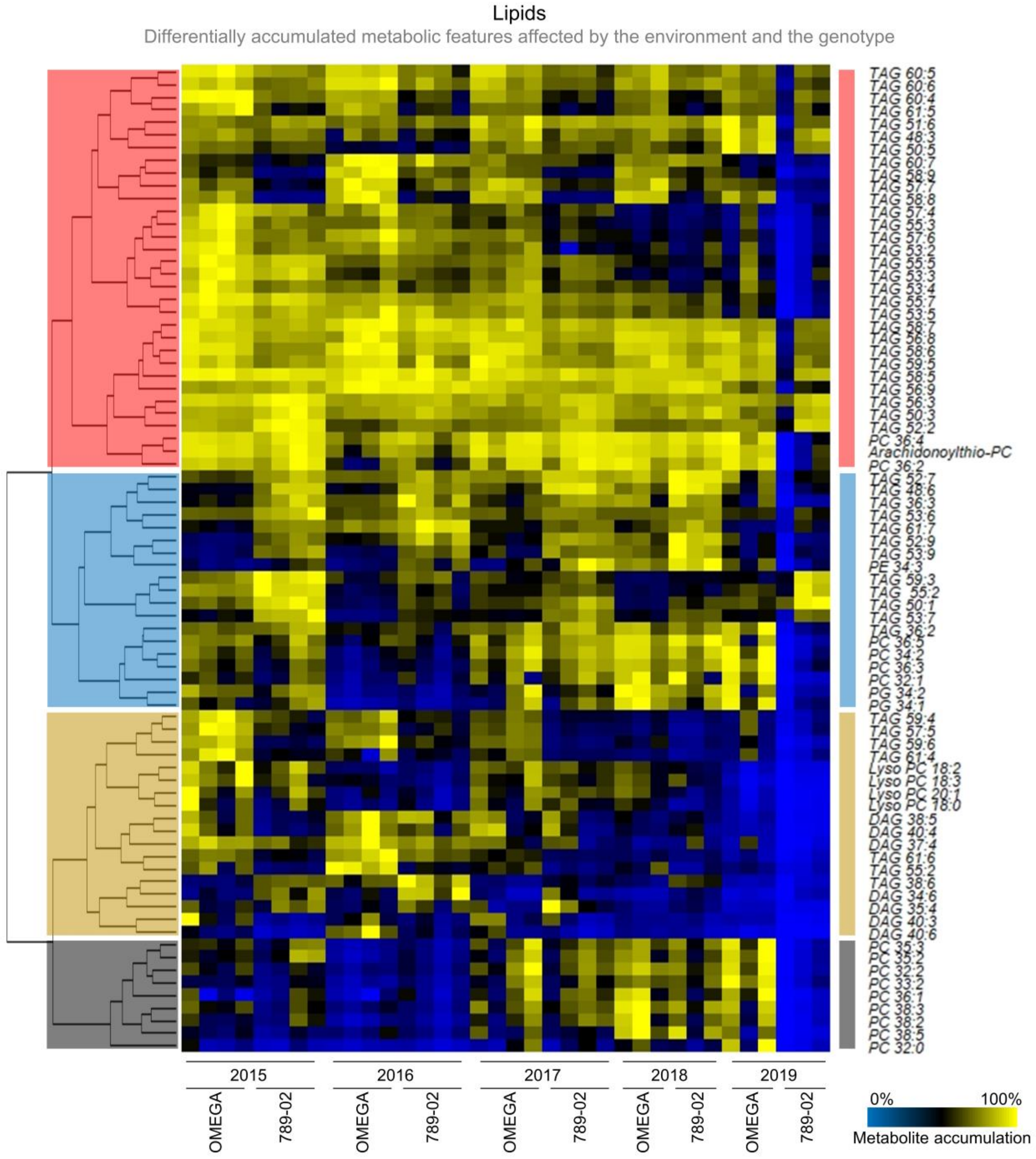


*Legend: DAG = Diacylglycerols; TAG = Triacylglycerols; PC = phosphatidylcholine; PE = phosphatidylethanolamine*
